# The ancillary N-terminal region of the yeast AP-1 transcription factor Yap8 contributes to its DNA binding specificity

**DOI:** 10.1101/614503

**Authors:** Ewa Maciaszczyk-Dziubinska, Anna Reymer, Nallani Vijay Kumar, Wojciech Białek, Katarzyna Mizio, Markus J. Tamás, Robert Wysocki

## Abstract

Activator protein 1 (AP-1) is one of the largest families of basic leucine zipper (bZIP) transcription factors in eukaryotic cells. How AP-1 proteins achieve target DNA binding specificity remains elusive. In *Saccharomyces cerevisiae*, the AP-1-like protein (Yap) family comprises eight members (Yap1 to Yap8) that display distinct genomic target sites despite high sequence homology of their DNA binding bZIP domains. In contrast to the other members of the Yap family, which preferentially bind to short (7-8 bp) DNA motifs, Yap8 binds to an unusually long DNA motif (13 bp). It has been unclear what determines this unique specificity of Yap8. In this work, we use molecular and biochemical analysis combined with computer-based structural design and molecular dynamics simulations of Yap8-DNA interactions to better understand the structural basis of DNA binding specificity determinants. We identify specific residues in the N-terminal tail preceding the basic region, which define stable association of Yap8 with the *ACR3* promoter. We propose that the N-terminal tail directly interacts with DNA and stabilizes Yap8 binding to the 13 bp motif. Thus, beside the core basic region, the adjacent N-terminal region contributes to alternative DNA binding selectivity within the AP-1 family.

## INTRODUCTION

Yap8 protein is one of eight members of the yeast AP-1 (Yap) family (1) that belongs to the fungal specific Pap1 subfamily of basic leucine zipper (bZIP) transcription factors (2) (Figure 1). The bZIP proteins regulate transcription by binding as dimers to specific DNA motifs. Yap1 preferentially binds to a 7 bp pseudo-palindromic sequence TTACTAA called the Yap response element (YRE) (1). However, Yap1 can also recognize TGACTAA (3,4), TGAGTAA (5) and TGACAAA (5) motifs. Other members of the Pap1 subfamily, like *Schizosaccharomyces pombe* Pap1, and *S. cerevisia*e Yap4 and Yap6, have preferences for an 8 bp palindromic version of the YRE (TTACGTAA) (2,6). DNA binding of bZIP transcription factors involves amino acids in the conserved basic region that precedes the leucine zipper region involved in dimerization. The crystal structure of the Pap1 bZIP domain bound to the 8 bp YRE revealed that five amino acid residues of the basic region make direct contacts with a TTAC half-site, and these residues constitute the signature DNA recognition NxxAQxxFR sequence (2). This motif is highly conserved among members of the Pap1 subfamily suggesting that Yap proteins share a common mechanism of DNA binding (Figure 1). Despite the high similarity of DNA binding regions and corresponding recognition elements of 7-8 bp, little is known how individual Yap proteins achieve their specificity of transcriptional regulation.

**Figure 1.**
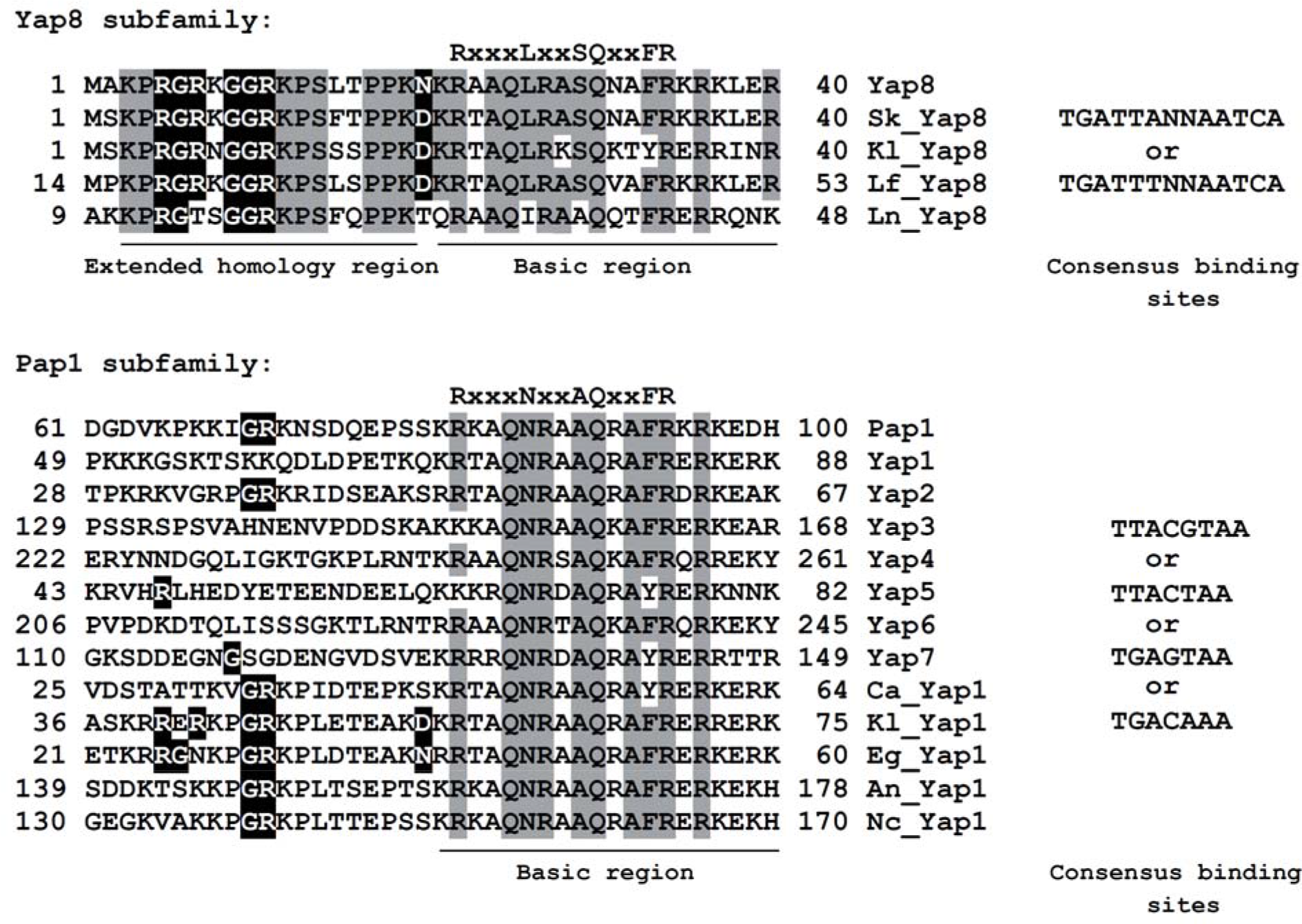
Comparison of basic regions, the N-terminal adjacent sequences and consensus DNA binding motifs of fungal AP-1 transcription factors. Conserved amino acid residues involved in direct binding to DNA bases as determined for the *Schizosaccharomyces pombe* Pap1 protein (NCBI accession no. NP_593662) (2) are indicated at the top of sequence alignment. Highly conserved residues in either Yap8 or Pap1 subfamily are in grey. N-terminal residues that are important for Yap8 binding to DNA are highlighted in black. Yap1-8 proteins are from *S. cerevisiae*, Ca_Yap1 is from *Candida albicans* (EEQ44283), Kl_Yap1 is from *Kluyveromyces lactis* (XP_451077), Eg_Yap1 is from *Eremothecium gossipii* (NP_984291), An_Yap1 is from *Aspergillus nidulans* (XP_680782), Nc_Yap1 is from *Neurospora crassa* (XP_957544), Sk_Yap8 is from *S. kudriavzevii* (EJT44313), Kl_Yap8 is from *Kluyveromyces lactis* (XP_453958), Lf_Yap8 is from *Lachancea fermentati* (SCW01455), Ln_Yap8 is from *L. nothofagi* (SCV05062). Consensus DNA binding motifs for each subfamily are indicated on the right panel.

The transcription factors Yap1 and Yap8 are key components of the cellular response to arsenite [As(III)], arsenate [As(V)] and antimonite [Sb(III)] stress. Yap1 and Yap8 sense the presence of these agents and coordinate activation of gene expression required for alleviation of metalloid toxicity (7–10). Yap1 stimulates transcription of a large set of genes encoding proteins that are involved in adaptation to arsenic-induced oxidative stress and metalloid detoxification (7,9,11,12). In contrast, Yap8 is highly specific and seems to activate transcription of only two genes (13); *ACR2* that encodes an arsenate reductase (14) and *ACR3* that encodes an As(III)/Sb(III) efflux transporter (15,16).

Yap8 is the only member of the Yap family that recognizes a long 13 bp TGATTAATAATCA sequence, called the Yap8 response element (Y8RE), that consists of a 7 bp core similar to the canonical YRE flanked by TGA bases (7,13). We recently showed that the Yap8 ortholog from *Kluyveromyces lactis* binds to multiple variants of Y8RE with different 7 bp core sequences flanked by conserved TGA bases (17). That study together with mutational analysis of the Y8RE sequence in *S. cerevisiae*, highlighted the importance of flanking TGA bases for Yap8-DNA interactions (Figure 1) (13,17). This distinct DNA binding property of Yap8 is reflected in its basic region in which invariant Asn and Ala residues of the NxxAQxxFR consensus sequence are replaced with Leu and Ser (LxxSQxxFR) (Figure 1). Indeed, Leu26 is essential for Yap8 binding to Y8RE and has, together with Asn31 and Leu26, been proposed to contribute to the DNA binding specificity of Yap8 (18).

In this study, we determined that a Yap8 variant with a core basic region identical to that of Yap1 still binds to Y8RE and fully activates transcription of *ACR3*. Such Yap8 variants also acquire capacity to bind to some, but not all, 7 bp motifs recognized by Yap1. Mutational analysis of the N-terminal tail adjacent to the basic region revealed specific residues that are required for stable association of Yap8 with the Y8RE-containing *ACR3* promoter and its activation. Based on a Yap8-DNA interaction model, we propose that the N-terminal tail directly interacts with DNA and stabilizes Yap8 binding to the 13 bp motif. We propose that the N-terminal tail of Yap8 constitutes an ancillary region that contributes to a unique DNA binding activity of Yap8 towards the 13 bp-long Y8RE motif. We hypothesize that the N-terminal region preceding the core basic region influence the DNA binding specificity of other AP-1 proteins.

## MATERIALS AND METHODS

### Strains, plasmids and growth conditions

The *S. cererevisiae* strains used in this study were wild type W303-1A (*MAT***a***ade2-1 can1-100 ura3-1 his3-11,15 leu2-3,112 trp1-1*), RW104 (*acr3Δ::kanMX*), RW117 (*yap8*Δ*::loxP*), RW120 (*yap8*Δ*::loxP yap1*Δ*::loxP::kanMX::loxP*), and RW124 (*yap1*Δ*::loxP*) (7). Plasmids used in this study are described in Supplementary Table S1. Standard yeast methods and growth conditions were used. Growth assays in the presence of sodium arsenite (Sigma-Aldrich) were carried out as previously described (19).

### Mutagenesis

Site-directed mutagenesis of *YAP8* was performed using pYX122-YAP8 (20) and pGEX4T-1-GST-YAP8 (13) plasmids as templates, the oligonucleotides listed in Supplemental Table S2 and QuikChange Lightning Site-Directed Mutagenesis Kit (Agilent Technologies) according to the protocol provided by the manufacturer. All mutations were confirmed by commercial DNA sequencing.

### β-galactosidase assay

Yeast cells expressing various versions of *ACR3-lacZ* gene fusions were grown in selective minimal medium in the presence of 0.1 mM As(III) for 6 h or left untreated. The β-galactosidase activity was measured at least three times in triplicates on permeabilized cells as described previously (21).

### RNA extraction and quantitative real-time PCR (qRT-PCR)

Total RNA was isolated from exponentially growing cells that were either untreated or exposed to 0.1 mM As(III) and collected at the indicated time points using RNeasyMini Kit (Qiagen). Reverse transcription was performed with 1.5 µg of purified RNA using High-Capacity cDNA Reverse Transcription Kit (Applied Biosystems) according to the manufacturer’s instruction. Quantitative real-time PCRs were performed in the LightCycler 480 Instrument (Roche), using RealTime 2xPCRMaster Mix SYBR (A&A Biotechnology) and ACR3-fw/rv primers listed in Supplemental Table S2 as described previously (22). *IPP1* was used as a reference gene. All assays were performed at least three times (biological replicas) in triplicates (technical replicas).

### Protein extraction and Western blot analysis

Cell extracts were prepared by TCA precipitation and proteins were separated by 10% SDS-PAGE followed by immunoblotting with anti-HA antibody (Sigma-Aldrich, ref: H6908, lot: 015M4868V, 1:2500 dilution) and anti-PGK1 antibodies (Abcam, ref: ab11368, lot: GR254438-1; 1:5000 dilution).

### Immunofluorescence microscopy

Immunofluorescent labeling of yeast cells was performed as described earlier (23). Cells were fixed in 3.7% formaldehyde for 2 h, washed and digested with Zymolyase (BioShop) for 30 min. The efficiency of spheroplasting was monitored by phase microscopy. Spheroplasts were washed twice and suspended in PBS buffer supplemented with 0.1% BSA. Yeast cells were stained with primary antibody (anti-HA, Sigma-Aldrich, ref: H6908, lot: 015M4868V, 1:1000 dilution) for 12 h at 4°C. The samples were washed with PBS containing 0.1% BSA after exposed secondary antibody Alexa Fluor® 488 goat anti-rabbit IgG (H+L, Life Technologies, ref: A11008, lot: 1470706, 1:200 dilution) at room temperature for two hours. After triple washing with PBS, cells were labeled with DAPI (Life Technologies, 1:5000 dilution) to visualize nuclei and examined with a fluorescence microscope (Axio Imager M2, Carl Zeiss) equipped with a 100x oil immersion objective, differential interference contrast and appropriate filters. Images were collected using Zeiss AxioCam MRm digital camera and processed with Zeiss Zen 2012 software.

### Expression and purification of GST-Yap8 variants

Expression of wild type and mutant versions of GST-Yap8 was induced by incubating *Escherichia coli* BL21(DE3)pLysS cells with 1 mM IPTG (isopropyl β-D-thiogalactoside) for four hours at 30°C in LB medium (1% tryptone, 0.5% yeast extract, 1% NaCl) in the presence of 100 μg/ml chloramphenicol. Cells were harvested and disrupted by sonication in cold PBS buffer containing protease inhibitor cocktail (Roche), 10 mM β-mercaptoethanol, 1% Triton X-100 and 10% glycerol. All GST-tagged proteins were purified using glutathione beads (GE Healthcare) according to the protocol supplied by the manufacturer.

### Electrophoretic mobility-shift assay (EMSA)

The 5’ end biotinylated complementary oligonucleotide pairs (Sigma-Aldrich) were annealed to make double-stranded and biotin-labeled probes by mixing in a buffer (10 mM Tris-HCl, pH 8.0, 1 mM EDTA), boiling for 5 min and cooling slowly to room temperature. Unlabeled complementary oligonucleotide pairs were also annealed to make double-stranded competitor probes. EMSA reaction solutions were prepared by adding the following components according to the manufacturer’s protocol (LightShift Chemiluminescent EMSA kit; Thermo Fisher Scientific): 1× binding buffer, 50 ng poly (dI-dC), 2.5% glycerol, 0.05% Nonidet P-40, 5 mM MgCl_2_, 10 ng of purified recombinant GST-tagged protein, competitor (4 pmol) and biotin-labeled probes (20 fmol). Reaction solutions were incubated for twenty minutes at room temperature. The protein-probe mixture was separated in a 6% polyacrylamide native gel in a standard 0.5× TBE buffer. Electrophoresis was performed on ice (100 V, 1 h). The DNA was transferred (100 V, 30 min) to a positive nylon membrane (Amersham Hybond-N+, GE Healthcare) and UV crosslinked (1200 uJ/cm2, UVP TL-2000 Ultraviolet Translinker). Migration of biotin-labeled probes was detected in the ChemiDoc MP Imager (BioRad) using streptavidin-horseradish peroxidase conjugates that bind to biotin and chemiluminescent substrate according to the manufacturer’s protocol. The sequences of the oligonucleotides used are listed in Supplementary Table S2.

Alternatively, the oligonucleotide probes were 5’ end labeled with [γ-^32^P]ATP using polynucleotide kinase (Thermo Scientific), purified through Sephadex G-50 chromatography, annealed with complementary oligonucleotides in the presence of 100 mM NaCl at 75°C for 10 min and gradually cooled to room temperature. Purified recombinant GST-tagged proteins (at indicated concentrations) were incubated with ^32^P-labeled oligonucleotide probes (40,000 cpm) in a 20 μ reaction containing EMSA buffer (10 mM Tris-HCl, pH 8.0, 50 mM NaCl, 1 mM DTT, 0.05% NP-40, 100 ng poly(dI-dC) and 6% glycerol) for 30 min at 4°C. The reaction mixtures were subjected to electrophoresis on 5% non-denaturing polyacrylamide gels in a standard 0.5x TBE buffer. Electrophoresis was performed on ice (100 V, 1 h). The gels were dried and analyzed using a phosphorimager (Molecular Imager FX, Bio-Rad).

### Fluorescence anisotropy assay

The fluorescence anisotropy of a FAM-labeled *ACR3* oligonucleotide (5’-CTTTTTGTTTGATTAATAATCAACTTTAGCG-3’, labeled on the 5’ end with 6-carboxyfluorescein) was measured on two-four independent repetitions with different protein to DNA ratios and one reference solution without protein in buffer A (10 mM Tris-HCl pH 8, 50 mM NaCl, 0.5% NP-40, 1 mM DTT and 5% glycerol). The total volume of the working solutions was 100 µl, and the added protein elution buffer amount was kept constant to 50 µl by adding buffer A when necessary. Measurements were performed on a spectrofluorometer FS5 (Edinburgh Instruments) in a temperature-controlled microcuvette at 25°C. Fluorescence emission intensity was recorded at 515 nm, with excitation at 490 nm, and emission and excitation slits set to 2 nm. All titrations were performed using 1 nM of DNA, and after each addition the sample was equilibrated for 6 min. Stoichiometric binding curves were fit to the equation: Δ*A*= Δ*A*_T_/2*D*_T_{(*E*_T_+*D*_T_+*K*_d_) – [(*E*_T_+*D*_T_+*K*_d_)^2^ – 4*E*_T_*+D*_T_]^1/2^}, where Δ*A* is the change in anisotropy, Δ*A*_T_ is the total anisotropy change, *E*_T_ is the total protein concentration, *D*_T_ is the total DNA concentration, and *K*_d_ is the dissociation constant.

### Chromatin immunoprecipitation (ChIP)

ChIP was performed as described previously (24). Sheared chromatin was immunoprecipitated using anti-HA antibody (Sigma-Aldrich, ref: H6908, lot: 015M4868V, 1:2500 dilution) overnight followed by incubation with sepharose protein G beads (Dynabeads Protein G, Life Technologies). Precipitates and input DNA were analyzed by qRT-PCR using PRACR3-fw/rv oligonucleotides listed in Supplementary Table S2. Fold enrichment represents the ratio of recovered DNA to input DNA of the *ACR3* promoter region from −251 to −100 relative to the ATG translation initiation codon normalized to the same ratio obtained for the *IPP1* gene. These ratios were additionally normalized to pre-induction (0 min) values and corrected for 0.5 mM As(III) induction (30 min).

### Molecular modeling

The model structure of the basic-leucine-zipper domain (residues 7-89) of the Yap8 protein homodimer was created using the homology building functionality of Yasara program (25). The DNA sequence of 25 bp (TTTGTT-**TGATTAATAATCA**-ACTTTA) contains Yap8-response element, Y8RE, shown in bold. The structure of the Yap8-DNA complex was modeled using HADDOCK molecular multi-body docking server (26,27). The residues: Asn20, Arg22, Gln25, Leu26, and Phe33 were indicated as "active", as their alanine-mutants show sufficient reduction in the proteins activity and/or ability to bind DNA (Table 1). Residues Arg27 and Arg36 were defined as "passive" as their alanine mutants show only partial resistance to As(III). The Y8RE-DNA residues were identified as "passive". Out of the 29 structure clusters provided by the HADDOCK server, one of the clusters had a significantly higher score, which was selected for further analysis. Lastly, the N-terminal fragments (residues 7-18) of the protein were added manually in a random coil configuration using program USCF Chimera (28). The random coil configuration of the N-terminal tails was justified by protein secondary structure prediction servers Jpred4 (29), PredictProtein (30) and PSIPRED (31). Additionally, the complex structure containing Asn20Ala Yap8 mutants was created using the ‘Rotamers’ functionality of USCF Chimera program.

**Table 1.** Summary of functional analysis of Yap8 mutant proteins.

| Mutant name | Mutated region | As(III) resistance | Mutant class | Model prediction |
| --- | --- | --- | --- | --- |
| P4A | N-term | +++ | F | Not included in the model |
| R5A | N-term | + | PF | Not included in the model |
| G6A | N-term | + | PF | Not included in the model |
| R7A | N-term | + | PF | interacts with DNA bases and backbone |
| K8A | N-term | +++ | F | Residue's backbone interacts with DNA backbone |
| G9A | N-term | + | PF | Plasticity of N-term region, tighter contact with DNA |
| G10A | N-term | – | NF | Plasticity of N-term region, tighter contact with DNA, interacts with DNA backbone |
| R11A | N-term | + | PF | Interacts with DNA backbone |
| K12A | N-term | +++ | F | Interacts with DNA backbone |
| P13A | N-term | +++ | F | Model shows no contact with DNA |
| S14A | N-term | +++ | F | Interacts with DNA backbone |
| L15A | N-term | +++ | F | Model shows no contact with DNA |
| T16A | N-term | +++ | F | Model shows no contact with DNA |
| P17A | N-term | +++ | F | Model shows no contact with DNA, caps the basic region |
| P18A | N-term | +++ | F | Model shows no contact with DNA, caps the basic region |
| K19A | N-term | +++ | F | Some contact with DNA backbone |
| N20A | N-term | + | PF | No contact with DNA, defines the conformational space of N-terminal region to enable tighter protein-DNA contacts |
| K21A | N-term | +++ | F | Interacts with DNA backbone, seen only for monomer 2 |
| R22A* | Basic | – | NF | Direct contact with <u>TGA</u> bases of Y8RE |
| A23 | Basic | ND | ND | Model shows no contact with DNA |
| A24 | Basic | ND | ND | Model shows no contact with DNA |
| Q25A | Basic | – | NF | Interacts with DNA backbone |
| L26A* | Basic | – | NF | Model shows no contact with DNA, but this observation is sensitive to definition of a hydrophobic interaction |
| R27A* | Basic | ++ | PF | Interacts with DNA backbone |
| A28 | Basic | ND | ND | Model shows no contact with DNA |
| S29A* | Basic | +++ | F | Interacts with DNA backbone |
| Q30 | Basic | ND | ND | Forms several H-bonds with bases from the central region of Y8RE |
| N31A* | Basic | +++ | F | Some contact with DNA backbone |
| A32 | Basic | ND | ND | Model shows no contact with DNA |
| F33 | Basic | – | NF | Hydrophobically interacts with <u>TGATT</u> |
| R34 | Basic | ND | ND | Interacts with DNA backbone and central A base of Y8RE |
| K35A | Basic | +++ | F | Model shows no contact with DNA |
| R36A | Basic | ++ | PF | Direct contact with DNA backbone |
| K37A | Basic | +++ | F | Model shows no contact with DNA |
| L38A | Basic | +++ | F | Model shows no contact with DNA |
| E39A | Basic | +++ | F | Model shows no contact with DNA |
| R40A | Basic | ++ | PF | Model shows no contact with DNA |

Mutated region: N-term refers to N-terminal region preceding the basic region. As(III) resistance: –, none; + or ++, partial; +++, full. Mutant class: F – functional, PF – partially functional, NF – non-functional. ND – not determined. \*Determined by Amaral *et al.*, Plos One (2013).

The two complex structures were subjected to subsequent studies by molecular dynamics simulations (MD), using GROMACS MD software package, version 5.1(32). Simulations were carried out using a combination of the latest AMBER all-atom nucleic acid Parmbsc1 (33) and ff14SB (34) force fields in implicit solvent using of SCP/E water molecules (35) and 150 mM KCl. MD simulations were carried out at constant pressure and temperature (1 atm., 300 K). Further details of the simulation protocols can be found in Supplemental Information. Each productive MD run was 500 ns long. MD trajectories were analyzed using CPPTRAJ program (36), focusing on the analysis of the protein-DNA interactions, including hydrogen bonds, salt bridges, and hydrophobic (apolar) interactions. Dynamic contacts maps were created by summing up the hydrogen bonds and the salt bridge interactions for each pair of Yap8-DNA interacting resides, which resulted in a contact strength value. We also performed conformational clusters analysis following the protocol described by Lavery and co-workers (37,38) for the basic-regions (residues 17-40) of the protein dimer. For the random-coil N-terminal regions (residues 7-16) conformational clusters were identified with the cluster feature of CPPTRAJ program (36), using DBSCAN (density-based) clustering algorithm (39). RMSD of heavy atoms of DNA outside YRE region and excluding two terminal base pairs on both ends and the protein residues 7-16 was used as a distance metric. The Yap8-DNA complex structure that represents the biggest conformational cluster was selected to represent the model structure. Molecular graphics were created with USCF Chimera.

## RESULTS

### Construction of Yap8 variants containing the Yap1-like basic region

The DNA binding basic region is highly conserved in the fungal AP-1 family (Figure 1 and 2A). Mutations of several conserved residues in the Yap8 basic region, including Arg22, Gln25, Arg27, and Arg36, were previously reported to impair the transcriptional activity of Yap8 towards a Y8RE-containing promoter (18). Also in our hands, Yap8-Q25A was not able to induce expression of the *ACR3-lacZ* reporter gene (Figure 2B) or rescue As(III) sensitivity of cells lacking the *YAP8* gene (Figure 2C). Likewise, Yap8-R36A appeared partially defective as we observed weak activation of *ACR3-lacZ* expression (Figure 2B) and partial complementation of *yap8*Δ (Figure 2C). However, Yap8 contains several amino acid substitutions within its basic region at positions conserved in other members of Yap family (Figure 1); these amino acid residues may contribute to the specificity of Yap8 towards the extended Y8RE motif as well as its inability to bind to short YRE motifs.

**Figure 2.**
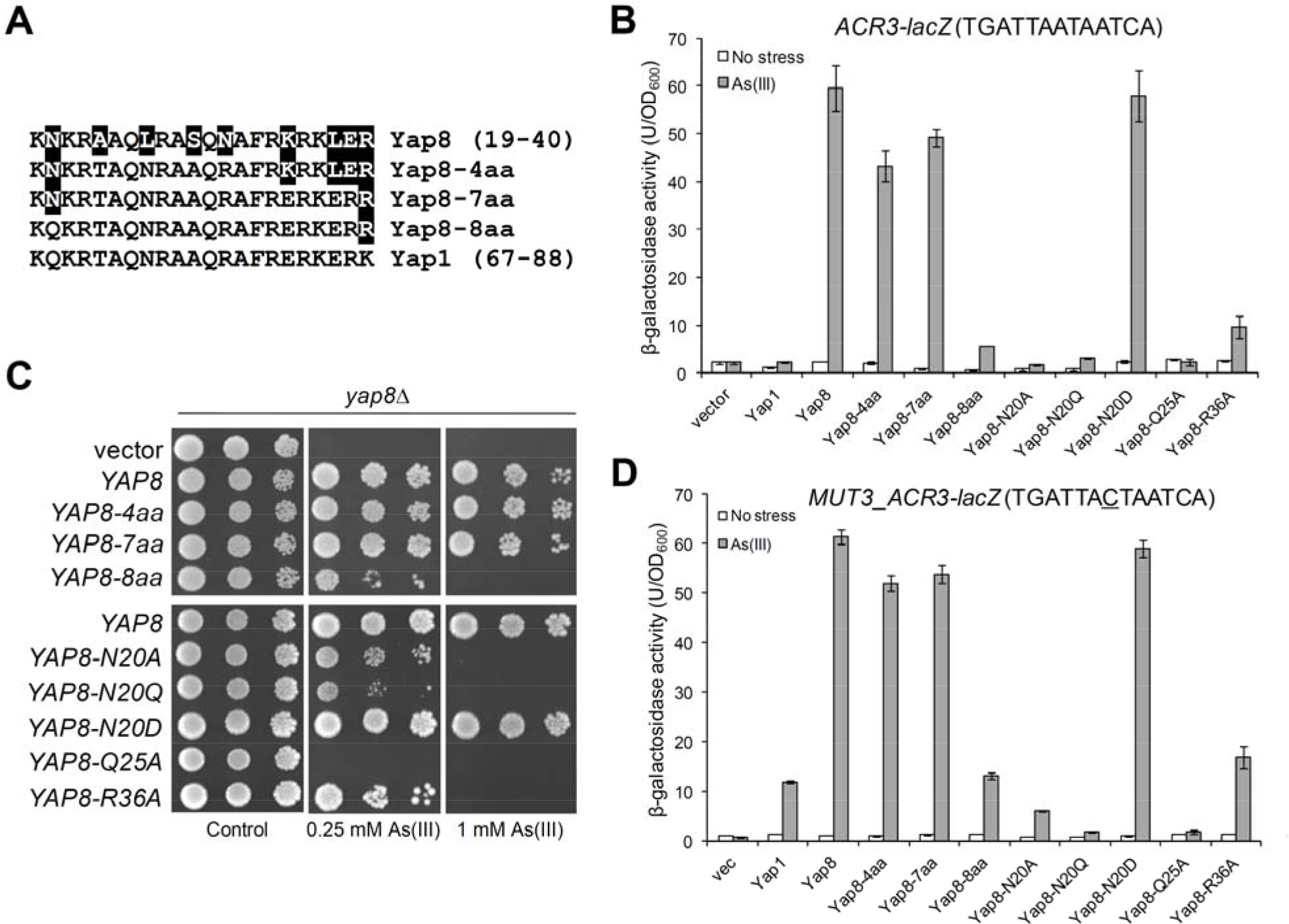
Analysis of Yap8 basic region variants. (**A**) Mutagenesis strategy to stepwise turn the Yap8 basic region into Yap1-like sequences. (**B**) β-galactosidase activity driven by the *ACR3-lacZ* promoter was measured in the *yap1*Δ *yap8*Δ mutant expressing either Yap1, Yap8 or Yap8 mutant proteins. Cells were exposed to 0.1 mM As(III) for 6 h or left untreated for the control. The values are the means of three biological replicas performed in triplicate ± S.D. (**C**) Complementation of As(III) sensitivity of *yap8*Δ by Yap8 variants. The *yap8*Δ mutant was transformed with empty vector (pYX122) or plasmids expressing indicated Yap8 variants. The resulting transformants were spotted on minimal selective plates containing various concentrations of As(III) and incubated 3 days at 28°C. (**D**) β-galactosidase activity driven by the *MUT3_ACR3-lacZ* promoter was measured in the *yap1Δ yap8*Δ mutant expressing either Yap1, Yap8 or Yap8 mutant proteins. Cells were exposed to 0.1 mM As(III) for 6 h or left untreated for the control. The values are the means of three biological replicas performed in triplicate ± S.D.

To investigate this, we stepwise replaced amino acid residues in the basic region of Yap8 into the corresponding residues present in Yap1 and functionally characterized the resulting Yap8 variants. First, we constructed a quadruple A23T L26N S29A N31R mutant (or Yap8-4aa) to make the core of the basic region identical with the Yap1 basic region, including the NxxAQxxFR consensus sequence (Figure 2A). The Yap8-4aa mutant was able to fully activate expression of the *ACR3*-*lacZ* reporter gene (Figure 2B) and to complement the As(III) sensitivity of the *yap8*Δ mutant (Figure 2C). Next, we introduced three additional mutations (K35E, L37E, and E39R) to make the C-terminal region adjacent to the core basic region identical to the Yap1 sequence (Figure 2A). The septuple Yap8-7aa mutant also behaved like wild type Yap8 in terms of *ACR3* expression and *yap8*Δ complementation (Figure 2B and C). Finally, we additionally replaced Asn20, located adjacent to the core of the basic region with Gln (corresponding amino acid in Yap1) in Yap8-7aa (Figure 2A). The octuple Yap8-8aa mutant failed to *trans*-activate the *ACR3*-*lacZ* reporter gene and complement As(III) sensitivity of *yap8*Δ (Figure 2B and C). In this regard, Yap8-8aa behaved like Yap1, which is not able to activate *ACR3* expression (Figure 2B). However, if the central adenine residue in the Y8RE element is replaced with cytosine, Yap1 can weakly induce *ACR3* expression (13) (Figure 2D). Thus, we analyzed activity of the *MUT3-ACR3-lacZ* promoter with the TGA<u>TAACTAA</u>TCA sequence containing both Y8RE and YRE (underlined) motifs in a single element (Figure 2D). Wild type Yap8, Yap8-4aa and Yap8-7aa variants strongly induced expression of the *MUT3-ACR3-lacZ* reporter gene whereas Yap8-8aa behaved like Yap1 and weakly activated the *MUT3-ACR3* promoter (Figure 2D). In sum, these results suggest that Asn20 contributes to Yap8 specificity towards the *ACR3* promoter.

To get a better insight into the role of Asn20, we replaced this residue with glutamine (Yap8-N20Q) or aspartate (Yap8-N20D), which are present in the corresponding positions in Yap1 and *K. lactis* Yap8, respectively (Figure 1). Asn20 was additionally replaced with alanine (Yap8-N20A). Yap8-N20A and Yap8-N20Q failed to induce expression of both *ACR3-lacZ* and *MUT3-ACR3-lacZ* upon As(III) stress (Figure 2B and D) and poorly complemented As(III) sensitivity of the *yap8*Δ mutant (Figure 2C). In contrast, Yap8-N20D showed wild type activity in both assays (Figure 2B and C).

We confirmed that all Yap8 variants tested were present at the same amounts as the wild type Yap8 protein (Supplementary Figure S1) and that all tested variant proteins were correctly localized to the nucleus (Supplementary Figure S2). Thus, the observed effects are likely directly related to Yap8 function/activity. We conclude that both the Yap8-specific (Asn20) residue adjacent to the basic region as well as highly conserved residues (Gln25, Arg36) within the basic region are important for Yap8 function.

### DNA binding properties of the Yap8 basic region and N20A mutants

To characterize the DNA binding properties of the Yap8 variants, we performed EMSAs using purified GST-Yap8 proteins (Supplementary Figure S3) and biotin-labeled oligonucleotides corresponding to the *ACR3* promoter sequence with the TGATTAATAATCA motif (Figure 3A). It is important to point out that we have previously shown that the GST-Yap8 fusion protein is fully functional *in vivo* (13). In agreement with published data (18) and our *in vivo* assays (Figure 2), Yap8-Q25A exhibited markedly reduced ability to bind to the *ACR3* oligo (Figure 3A). In line with the inability to induce *ACR3* expression (Figure 2C), Yap8-8aa and Yap8-N20A variants also showed highly reduced capacity to bind to the *ACR3* oligo (Figure 3A). Accordingly, *in vivo* ChIP experiment revealed that Yap8-8aa and Yap8-N20A do not stably associate with the *ACR3* promoter in living cells, neither in the absence nor presence of As(III) (Figure 3B). Importantly, the Yap8-7aa variant with Yap1-like core basic region retained the wild type activity both *in vitro* (Figure 3A) and *in vivo* (Figure 3B).

**Figure 3.**
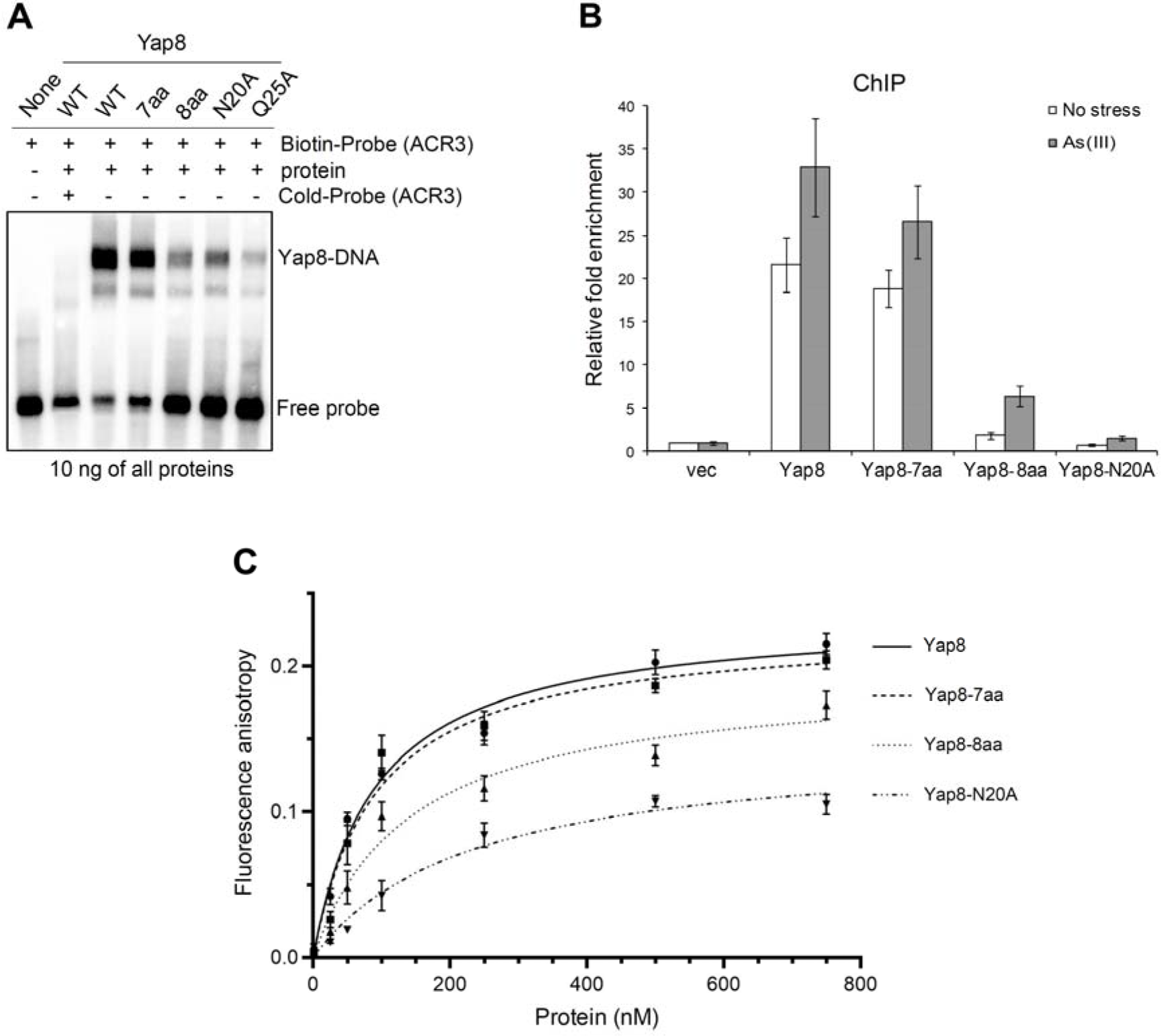
DNA binding activity of Yap8 basic region mutants. (**A**) Binding of Yap8 variants to the *ACR3* promoter as determined by EMSA. Purified GST-Yap8 variants at indicated concentration were incubated with biotin-labeled oligonucleotides corresponding to Y8RE-containing promoter fragments of *ACR3* gene followed by electrophoresis. (**B**) Binding of Yap8 variants to the *ACR3* promoter as determined by ChIP. *yap8*Δ cells bearing plasmids expressing the indicated Yap8-HA fusion proteins or the control vector were exposed to 0.5 mM As(III) for 30 min or left untreated. qRT-PCR was performed with chromatin fragments immunoprecipitated with anti-HA antibodies and primers amplifying the *ACR3* promoter region containing Y8RE1 and Y8RE2 motifs. Error bars indicate standard error of the mean from at least two independent biological replicas and four PCR reactions. (**C**) Fluorescence anisotropy assays performed with indicated variants of purified GST-Yap8 and the FAM-labeled *ACR3* promoter fragment as described in Materials and Methods.

To more accurately measure the affinity of Yap8-DNA interactions, we performed a fluorescence anisotropy binding assay (Figure 3C). Binding titrations were performed as a function of increasing concentration of Yap8 and its mutated forms at fixed DNA concentration corresponding to the *ACR3* promoter sequence. From fluorescence anisotropy measurements it is clear that the Yap8-7aa mutant showed virtually identical binding affinity as the wild-type protein. The values of *K*_*d*_ determined for the wild type Yap8 and the Yap8-7aa mutant version are 9.9 (±1.4) nM and 9.3 (±1.7) nM, respectively. In contrast, Yap8-8aa and Yap8-N20A variants showed significantly weaker binding to a fluorescein (FAM)-labeled Y8RE-containing DNA fragment with *K*_d_ values of 15.1 (±2.1) nM and 25.5 (±4.3), respectively. The results obtained by this solution-based, true-equilibrium method are consistent with *in vivo* (complementation tests, lacZ assay, ChIP) and *in vitro* (EMSA) data shown above (Figure 2B-D, Figure 3A and B). Together, our data strongly suggest that the N-terminal Asn20 residue is important for high affinity binding of Yap8 to the 13 bp long Y8RE motif.

### Yap8 variants with the Yap1-like basic region cannot bind to all YREs

We next investigated whether the Yap8 variants with Yap1-like basic regions had acquired ability to bind to 7 bp YRE motifs. For this, we performed EMSAs using oligos corresponding to *GSH1* (contains one YRE with sequence TTAGTCA) and *TRX2* (contains two YREs with sequence TTACTAA) promoters. None of these YREs contain TGA flanks. As expected, wild type Yap8 did not bind to the *GSH1* oligo (Figure 4A). However, Yap8-4aa, Yap8-7aa, and Yap8-8aa bound weakly to the *GSH1* oligo at higher protein concentrations (stable binding required 100 ng protein for the *GSH1* oligo compared to 10 ng for the *ACR3* oligo), suggesting low-affinity binding of these Yap8 variants to the YRE TTAGTCA (Figure 4). Neither Yap8-N20A nor Yap8-N20Q bound to the *GSH1* oligo. None of the Yap8 variants bound stably to the *TRX2* promoter fragment (Figure 4). Thus, replacing up to eight amino acid residues to make the Yap8 basic region more Yap1-like was not sufficient to enable binding of the modified Yap8 to the YRE motif (TTACTAA), present in *TRX2*. This suggests that the amino acid residues outside the basic region may contribute to YRE recognition and/or stable DNA binding.

**Figure 4.**
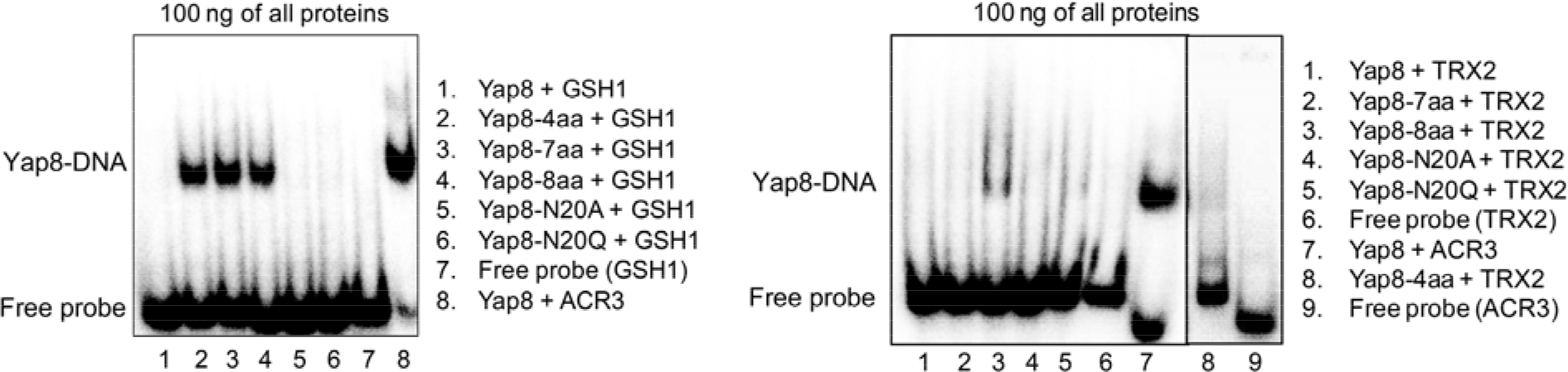
DNA binding of Yap8 protein variants with the Yap1-like basic region determined by EMSA. Purified GST-Yap8 variants at indicated concentrations were incubated with ^32^P-labelled oligonucleotides corresponding to Y8RE/YRE-containing promoter fragments of *ACR3*, *GSH1*, and *TRX2* genes followed by electrophoresis.

### The N-terminal tail adjacent to the basic region contributes to Yap8 DNA binding activity

The N-terminus of the bZIP domain of Yap1 (Asp63-Pro64) and Yap8 (Thr16-Pro17-Pro18) (Figure 1) include the N-capping motifs containing Asn, Asp, Ser, Thr, or Gly followed by single or double Pro residues. The N-capping motif is believed to stabilize the helical structure of the basic region upon DNA binding, without a direct interaction with DNA (40). To investigate the role of the putative N-cap of Yap8, we constructed and functionally characterized Yap8-T16A, Yap8-P17A, and Yap8-P18A mutants. We found that all tested mutants fully complemented the arsenic sensitivity of *yap8*Δ suggesting that the Thr-Pro-Pro motif does not affect the Yap8 function (Supplementary Figure S4). In addition, we tested the significance of adjacent Ser14, Leu15, Lys19 and Lys21 residues for Yap8 function by alanine replacement and found that the resulting mutants showed wild type phenotype (Table 1, Supplementary Figure S4). To summarize, it seems that in the Yap8 region of Ser14-Lys21 only Asn20 residue is important for Yap8 function.

Members of the mammalian Maf subfamily of bZIP superfamily that recognize a 13-14 bp consensus element (TGCTGAC(G)TCAGCA) called the Maf recognition element (MARE) (41) require the N-terminal extended homology region (EHR) preceding the basic region for high-affinity binding to DNA (42,43). Thus, we investigated whether the Pro4-Pro13 region of Yap8, which is rich in basic residues and highly conserved in the Yap8 subfamily (Figure 1), contributes to high-affinity binding to DNA. First, we constructed a Yap8 variant lacking residues from Arg5 to Pro13; the resulting Yap8-Δ5-13 mutant failed to complement arsenic sensitivity of *yap8*Δ strain (Figure 5A) and to induce expression of the *ACR3*-*lacZ* reporter gene (Figure 5B). Importantly, the Yap8-Δ5-13 mutant was expressed at the wild type level (Supplementary Figure S1) and showed nuclear localization (Supplementary Figure S2). This suggests that the N-terminal tail is critical for Yap8 function.

**Figure 5.**
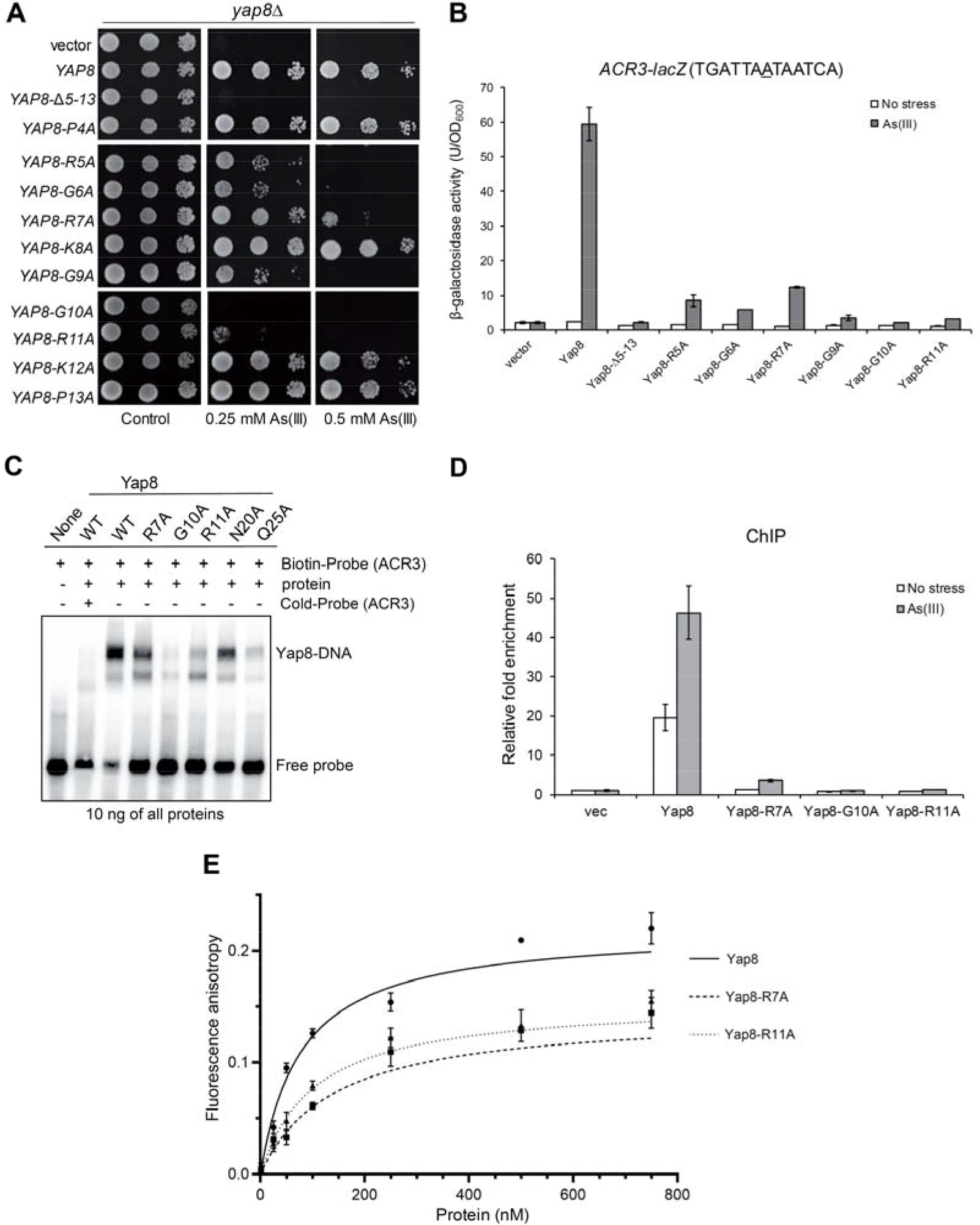
Functional analysis of Yap8 N-terminal tail mutants. (**A**) Complementation of As(III) sensitivity of *yap8*Δ by indicated Yap8 variants. The *yap8*Δ mutant was transformed with empty vector (pYX122) or plasmids expressing Yap8 variants. The resulting transformants were spotted on minimal selective plates containing various concentrations of As(III) and incubated 3 days at 28°C. (**B**) β-galactosidase activity driven by the *ACR3-lacZ* promoter was measured in the *yap1*Δ *yap8*Δ mutant expressing indicated Yap8 mutant proteins. Cells were exposed to 0.1 mM As ΔI) for 6 h or left untreated for the control. The values are the means of three biological replicas performed in triplicate ± S.D. (**C**) Binding of Yap8 variants to the *ACR3* promoter as determined by EMSA. Purified GST-Yap8 variants at indicated concentration were incubated with biotin-labeled oligonucleotides corresponding to Y8RE-containing promoter fragments of *ACR3* gene followed by electrophoresis. (**D**) Binding of Yap8 variants to the *ACR3* promoter as determined by ChIP. *yap8*Δ cells bearing plasmids expressing Yap8-HA variant proteins or the control vector were exposed to 0.5 mM As(III) for 30 min or left untreated. qRT-PCR was performed with chromatin fragments immunoprecipitated with anti-HA antibodies and primers amplifying the *ACR3* promoter region containing Y8RE1 and Y8RE2 motifs. Error bars indicate standard error of the mean from at least two independent biological replicas and four PCR reactions. (**E**) Fluorescence anisotropy assays performed with indicated variants of purified GST-Yap8 and the FAM-labeled *ACR3* promoter fragment as described in Materials and Methods.

To identify residues in the N-terminal tail that are important for Yap8 binding to DNA, we generated ten single alanine-replacement mutations covering residues from Pro4 to Pro13 and tested the functionality of the resulting Yap8 mutants. Of these: Yap8-R5A, Yap8-G6A, Yap8-R7A, and Yap8-G9A partially complemented arsenic sensitivity of *yap8*Δ strain (Figure 5A) and showed residual ability to induce expression of the *ACR3*-*lacZ* reporter gene (Figure 5B). Yap8-G10A and Yap8-R11A exhibited the strongest phenotype with no ability to confer resistance to As(III) (Figure 5A) or to activate the *ACR3* promoter (Figure 5B). Except for Yap8-G9A, which showed reduced protein level, all tested Yap8 mutants showed protein accumulation at the wild type level in response to As(III) treatment (Supplementary Figure S1) and nuclear localization both in the absence and presence of As(III) (Supplementary Figure S2). Based on these results, we conclude that the arginine and glycine-rich N-terminal region is important for Yap8 ability to activate the *ACR3* promoter.

We hypothesized that arginine residues of the Yap8 N-terminal tail may be involved in DNA binding, whereas glycine residues may contribute to plasticity of this region allowing tighter contact with DNA. To test this, we investigated the ability of purified Yap8-R7A, Yap8-G10A and Yap8-R11A protein variants tagged with GST (Supplementary Figure S3) to bind the *ACR3* promoter *in vitro* by EMSA (Figure 5C) and *in vivo* by ChIP (Figure 5D). Yap8-G10A showed no binding to DNA fragment containing the Y8RE motif, whereas Yap8-R7A and Yap8-R11A exhibited reduced binding to DNA compared to the wild type Yap8 (Figure 5C). Likewise, little (Yap8-R7A) or no association (Yap8-G10A and Yap8-R11A) of Yap8 variants to the *ACR3* promoter was observed by ChIP (Figure 5D). Next, we performed a fluorescence anisotropy binding assay to measure binding affinity of these Yap8 mutants to a DNA fragment comprising the *ACR3* promoter (Figure 5E). Both Yap8-R7A and Yap8-R11A mutant proteins showed decreased binding affinity to the Y8RE-containing DNA fragment. The values of *K*_d_ determined for the Yap8-R7A and Yap8-R11A variants were 13.4 (±1.7) nM and 18.9 (±2.9) nM, respectively, compared to *K*_d_ of 9.9 (±1.4) nM for the wild type Yap8. However, the affinity of Yap8-R7A and Yap8-R11A variants for the Y8RE-containing DNA fragment was stronger than the affinity observed for Yap8-N20A (25.5 ±4.3 nM) (Figure 3C). We were not able to perform this experiment for Yap8-G10A because of its high tendency of the purified protein to precipitate. In sum, based on these results we conclude that the N-terminal tail contributes to stable binding of Yap8 to DNA and a unique specificity of Yap8 towards the long (13 bp) Y8RE motif.

### Molecular modeling and structural analysis of Yap8-DNA complex

To get an atom-level understanding of the structural basis of Yap8-DNA recognition we created an all-atom model of Yap8-DNA complex (Figure 6). The model consists of Yap8 homodimer and 25 bp DNA segment containing the Y8RE motif. Each Yap8 monomer contains an α-helical basic-leucine-zipper (residues 17-89) domain and an unstructured N-terminal region (residues 7-16). Details of the model building are provided in the methods section. According to the model, the protein inserts its α-helical basic-leucine-zipper domain in the DNA major groove of the Y8RE sequence, with the N-terminal regions making contacts with the DNA minor groove of the Y8RE flanks. To verify the model we performed 500 ns all-atom molecular dynamics (MD) simulations. The MD simulation allowed construction of the dynamic interactions maps (Figure 7A), which describe the details of the specific Yap8-DNA interactions and the dynamics of the intermolecular interface. In addition, the protein-DNA interactions were characterized by the occupancy (percentage present) during the MD simulation and the average lifetime (Supplementary Table S3). The MD simulations show that the interactions patterns differ between the monomers (Figure 7A, Supplementary Table S3).

**Figure 6.**
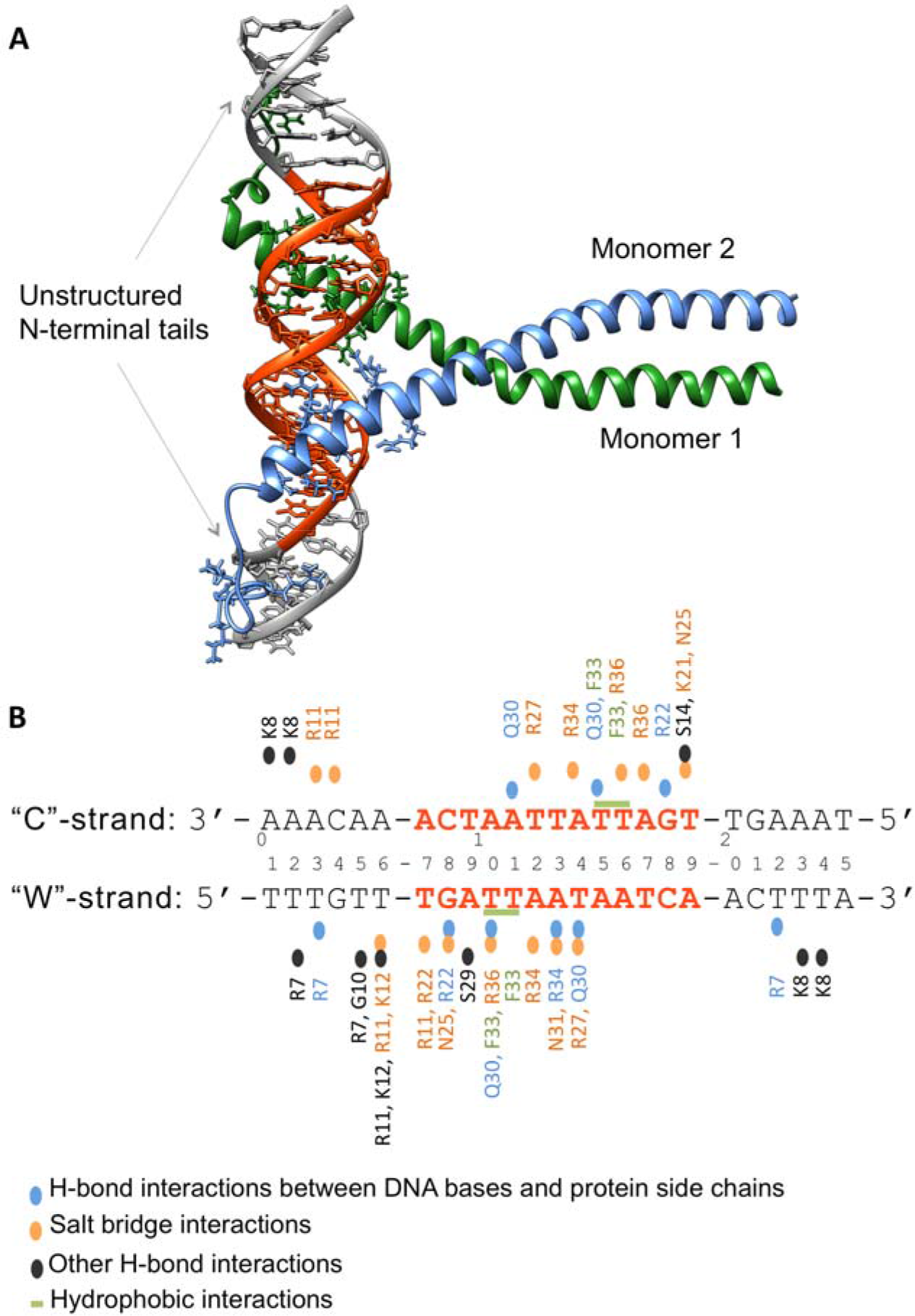
Model structure and interactions network of the Yap8-DNA complex. (**A**) Model structure of Yap8 protein homodimer in complex with the 25 bp long DNA segment containing Yap8 response element (Y8RE), in orange. Each Yap8 monomer includes an unstructured N-terminal region (residues 7-16) and basic leucine zipper domain (residues 17-89). (**B**) Schematic overview of the protein-DNA interactions. The DNA sequence, used in the model, is numbered 1-25 with the ‘Watson’ (“W”) strand representing the 5’-3’ direction and the ‘Crick’ (“C”) strand – the 3’-5’ direction. Only the interactions that occur at least 25% of the time of the 0.5 μ MD simulation are depicted.

**Figure 7.**
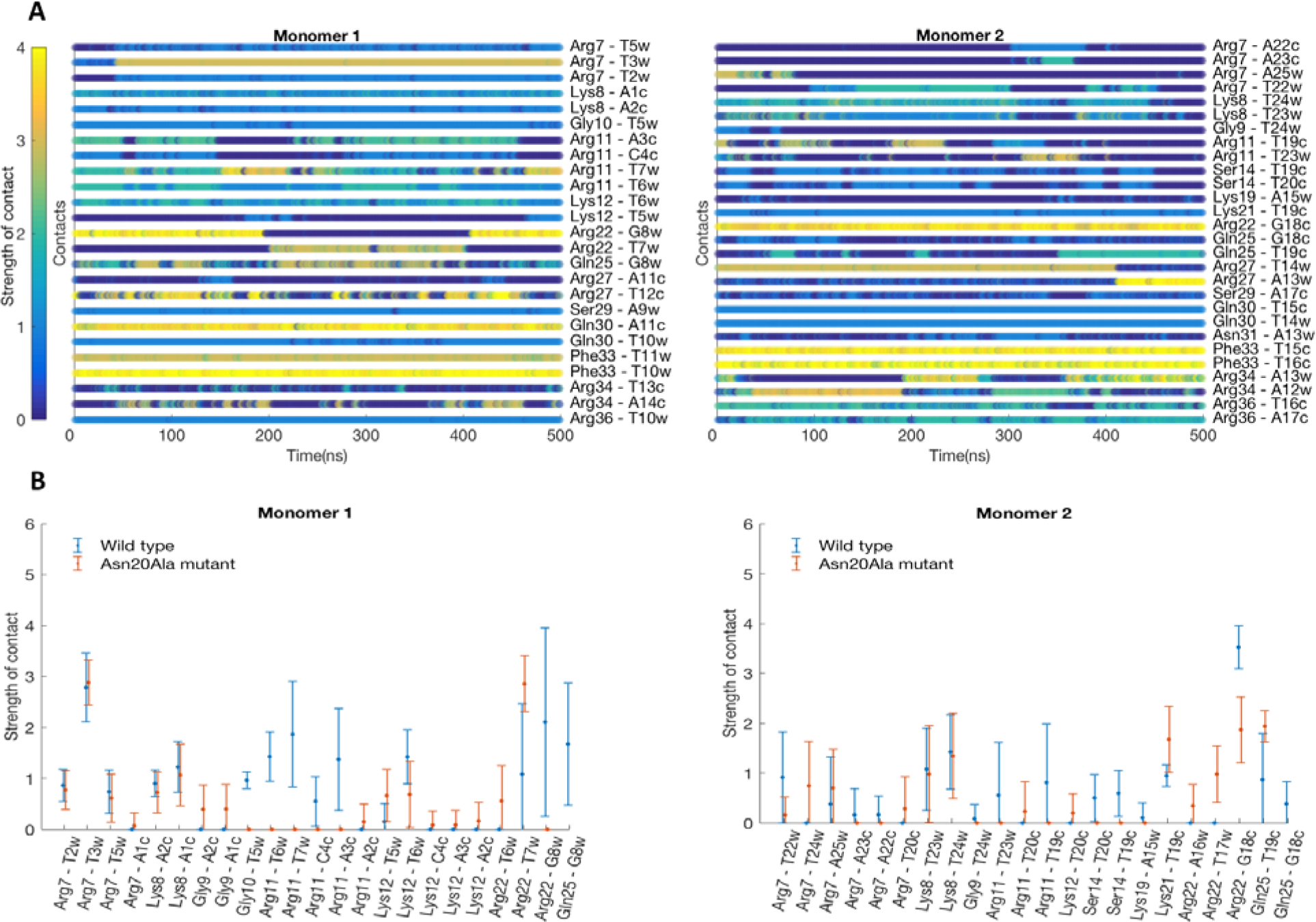
Characterization of Yap8-DNA interactions derived from the MD simulations. (**A**) Dynamic interactions maps illustrating the intermolecular wild-type Yap8-DNA interface. The interactions between pairs of the protein-DNA residues are characterized by a contact strength and its occurrence during the 0.5 μ MD simulation. (**B**) Comparison of the interaction patterns between pairs of residues of Yap8 wild-type or N20A-mutant proteins and DNA observed during the 0.5 μ MD simulations. Each specific contact is characterized by mean value of the contact strength and its standard deviation.

In the unstructured N-terminal regions (residues 7-16), we observe that Arg7 of both monomers form strong and stable hydrogen bonds with the T3_W_ and T22_W_ bases of the flanking sequences (subscripts "W" and "C" indicate correspondingly the 5’-3’ and the 3’-5’ DNA strands); Arg11 residues form salt bridges with the DNA backbone. But the occupancies and the lifetimes of the contacts vary between the monomers, presumably reflecting the different nucleotide composition of the Y8RE flanking sequences. There is also a number of hydrogen bonds formed between the backbones of the protein and DNA, which stabilize the N-terminals–DNA interactions, involving Arg7, Lys8, Gly10, Arg11 and Lys12. In the basic region (residues 17-40), we observe Arg22 and Asn25 residues interacting with the T7_W_ and G8_W_ bases of the TGA-triplet of the Y8RE-sequence. The model structure suggests that Leu26 of monomer 1 could participate in hydrophobic interactions with the methyl groups of T10_W_, and Leu26 of monomer 2 – with either T14_W_ or T15_C_ DNA bases. However, we do not observe these interactions during the MD simulations, although this observation is sensitive to the definition of a hydrophobic interaction. Here, we employed the 6Å distance between centers of masses of the corresponding residues. Gln30 of monomer 1 forms hydrogen bonds with the T10_W_ and A11_C_ bases, while Gln30 of monomer 2 forms hydrogen bonds with the T14_W_ and T15_C_ bases. Arg34 residue of monomer 2 forms a hydrogen bond with the A13W base. Arg34 of monomer 1 does not exhibit a symmetric interaction and only participates in a number of salt bridge contacts with the DNA backbone. Salt bridge contacts with the DNA backbone are also observed for Arg22, Asn25, Arg27, and Arg34 residues of monomer 1; and for Lys21, Asn25, Arg27, Asn31, Arg34, and Arg36 residues of monomer 2, though again the occupancies and the average lifetimes of the interactions differ between the monomers (Supplementary Table S3). Overall, monomer 2 appears to have a tighter interaction interface with the lower half of the YRE sequence (Figure 7A, Supplementary Table S3).

To investigate the role of Asn20 for the protein-DNA complexation, we repeated the 500 ns MD simulations for the N20A Yap8 mutant. Except for the N to A mutation in Yap8 N20A, the starting structures of the wild-type and the mutant complexes were identical. The intermolecular interface was again characterized by the dynamic interactions maps (Supplementary Figure S5), the contacts occupancies and average lifetimes (Supplementary Table S4). When comparing the wild-type Yap8-DNA and the N20A mutant-DNA interaction patterns, we observe a number of deviations in the N-terminal regions right before the start of the basic domain, residues 9-16. Interestingly the interactions with DNA exhibited by the residues further towards the N-terminus, Arg7 and Lys8, of both proteins are nearly identical in their strength, occupancy, and average lifetimes (Figure 7B). This observation suggests that Asn20 residue, even though not directly interacting with DNA might influence the conformational space of the N-terminal tails enabling a tighter protein-DNA contacts network.

## DISCUSSION

How does Yap8 achieve binding specificity towards its 13 bp recognition element? The characteristic feature of the Pap1 subfamily of bZIP proteins, including Yap1 to Yap7, is the presence of the conserved RxxxNxxAQxxFR motif in the DNA binding basic region (Figure 1). It has been shown for the Pap1-DNA complex that the signature residues Asn86, Ala89, Gln90, Phe93, and Arg94 are involved in direct interactions with DNA bases of the TTAC half-site of the 8 bp YRE whereas Arg82 binds to the guanine flanking the TTAC sequence (2). Four additional conserved Pap1 residues (Gln85, Arg87, Arg91, and Arg96) interact with the phosphate backbone (2). In Yap8, the conserved Asn and Ala residues in the DNA recognition sequence (<u>N</u>xx<u>A</u>QxxFR) are replaced with Leu26 and Ser29, and Arg91 involved in interaction with phosphate in Pap1 is replaced with Asn31 (Figure 1). Recently, alanine replacement analysis within the Yap8 basic region revealed the importance of the highly conserved residues Arg22, Gln25, Arg27, and Arg36 (corresponding to Arg82, Gln85, Arg87, and Arg96 of Pap1) for Yap8-DNA binding (Table 1) (18). In the case of the Yap8-specific residues Leu26, Ser29, and Asn31 (corresponding to Asn86, Ala89, and Arg91 of Pap1), only the L26A mutation impaired the DNA binding activity of Yap8. Interestingly, concomitant replacement of Leu26 and Asn31 with Asn and Arg (present in the corresponding positions in Pap1 and Yap1) extended the binding ability of Yap8 to the YRE motif (TTACTAA) as shown by *in vitro* EMSA assay (18). At the same time, the double L26N N31R Yap8 variant retained its ability to bind to Y8RE and complemented the arsenic sensitivity of the *yap8*Δ mutant (18). Consistent with these findings, we showed that Gln25 and Arg36 are important for Yap8 activity. Moreover, the quadruple A23T L26N S29A N31R mutant (or Yap8-4aa), having the core of the basic region identical with that of Yap1, retained full ability to induce *ACR3* expression and to bind Y8RE *in vitro* (Figure 2 and 3). Yap8-4aa showed low-affinity binding to the *GSH1* oligo containing the TTAGTCA motif (Figure 4) but was unable to bind to the *TRX2* promoter with two YREs (TTACTAA) (Figure 4). This suggests that structural elements outside the basic region may contribute to the DNA binding specificity of Yap proteins.

It has been previously shown that amino acid residues flanking the basic region are important for DNA-binding activity and DNA-target specificity of bZIP proteins (40,41,44). For example, transcription factors belonging to the mammalian Maf subfamily, bind to a 13-14 bp MARE consensus element (TGCTGAC(G)TCAGCA) (41,42). MARE consists of TGC and GCA flanks and the core motif TGACTCA (12-*o*-tetradecanoylphorbol 13-acetate (TPA)-responsive element, TRE) or TGACGTCA (cyclic AMP-responsive element, CRE). The CRE motif is also recognized by mammalian AP-1 (Jun-Fos heterodimer) and CRE binding protein (CREB/ATF), respectively. It was proposed that the N-terminal extended homology region (EHR) preceding the basic region (42) together with the substitution of the basic region Ala – a highly conserved residue, critical for DNA recognition in other AP-1 proteins (12) – with Tyr (RxxxNxx<u>Y</u>AxxCR) (45) determines the atypical binding specificity of Maf proteins. Unexpectedly, the X-ray crystal structure of the MafG-DNA complex revealed that the MafG-specific Tyr64 and EHR are not involved in MARE recognition (43). Instead, the invariant Arg57 and Asn61 residues (<u>R</u>xxx<u>N</u>xxYAxxCR), corresponding to Arg82 and Asn86 of Pap1, or Arg22 and Leu26 of Yap8 (Figure 1) directly contact the GC bases of the flanks instead of MARE. Binding of the basic region helix is stabilized by a network of hydrogen bonds formed by the residues of the basic region, including MafG-specific Tyr64, and several N-terminal residues either adjacent to the basic region or those forming short α-helices of EHR (43). Interestingly, the yeast bZIP transcription factor Hac1 involved in the unfolded protein response exhibits dual DNA binding specificity, and recognizes either short (6-7 bp) or extended (11-13 bp) motifs within target gene promoters (46). Importantly, the N-terminal region of Hac1 is required for the dual site recognition: the individual basic residues within this region contribute to the alternative specificities (46). To summarize, these observations suggest that N-terminal regions preceding the bZIP domains facilitate DNA binding and contribute to target gene specificity.

Here, we show that the N-terminal region adjacent to the basic region is critical for high affinity binding of Yap8 to the long (13 bp) Y8RE motif (Figure 3 and 5) and for induction of *ACR3* expression *in vivo* (Figure 2 and 5; Table 1). Mutational analysis of the Yap8 EHR revealed several arginines (Arg5, Arg7, Arg11) and glycines (Gly6, Gly9, Gly10), which are important for Yap8-dependent *ACR3* activation (Figure 5B). Moreover, we confirmed that Arg7, Gly10 and Arg11 facilitate Yap8 high affinity binding to the Y8RE motif (Figure 5C-E). The model of Yap8-DNA complex suggests that the N-terminal tail is engaged in a tight network of contacts between the protein and the Y8RE-DNA flanking sequences (Figure 7A), which enables stable positioning of the α-helical basic region in the major grooves of the Y8RE motif. The binding pose of the α-helical basic region allows contacts between Arg22, Asn25, Arg27, Ser29, Gln30, Phe33 and Arg36 residues and the extended TGATT half-site, while the central adenine base is recognized by Arg34 of Yap8 monomer 2. Interestingly, the contacts occupancies and average lifetimes, observed in 0.5 µs MD simulation, differ between the two monomers. This observation could result from the asymmetry of bZIP coil-coil protein dimerization, or the varied interactions patterns of the Y8RE-DNA flanks and the N-terminal regions.

Our data show that Asn20 adjacent to the basic region is critical for high affinity binding of Yap8 to the 13 bp long Y8RE motif. The model suggests that Asn20 is not in direct contact with DNA, but influences the conformational space of the N-terminal tails (region 9-19). The MD simulations of N20A Yap8 mutant-DNA complex showed that the mutant exhibits less stable contacts between the N-terminal and DNA, which could influence the overall stability of Yap8-DNA complexation.

Alignment of fungal AP-1 protein sequences revealed that residues corresponding to Gly10 and Arg11 in Yap8 are conserved in several member-proteins, including Pap1 (Figure 1). Of these, *K. lactis* Yap1 shows the closest similarity to the N-terminal tail of Yap8 and contains five of seven residues found to be important for Yap8 binding to the 13 bp Y8RE motif (Figure 1). We have previously shown that KlYap1 contributes to activation of *ACR2* and *ACR3* genes in *K. lactis* suggesting that it exhibits broader DNA binding specificity (17). Importantly, *K. lactis* Yap1, which contains the N-terminal region similar to Yap8, is able to partially complement lack of Yap8 in *S*. *cerevisiae* (our unpublished data). We propose that the composition of the N-terminal region preceding the basic region influences the repertoire of DNA motifs recognized by AP-1 proteins and dictate target gene specificities. It is important to emphasize that the MafG-DNA complex is the only crystal structure of bZIP domain dimer bound to DNA obtained with the protein fragment containing the N-terminal region (43). Investigating the significance of the N-terminal region of other bZIP proteins for DNA binding specificity will unveil the mechanisms employed by bZIP transcription factors for recognition of target gene sites.

## ACKNOWLEDGMENTS

We thank Beata Zagorska-Marek and Edyta Gola for providing antibodies for immunofluorescence and Scott Moye-Rowley for plasmids. The authors thank the Swedish National Infrastructure for Computing (SNIC) for the generous provision of computing resources. We also thank Prof. Leif A. Eriksson for many helpful discussions.

## Author contributions

E.M.D. designed and performed experiments and contributed to writing the manuscript; A.R. did the modeling of protein-DNA interactions and contributed to writing the manuscript; N.V.K. and K.M. performed experiments; W.B. designed and performed the anisotropy experiments; M.J.T. and R.W. contributed to the design of experiments and writing of the manuscript; all authors analyzed the results and approved the final version of the manuscript.

## FUNDING

This work was supported by grants 2012/07/B/NZ1/02804 (to R.W.) and 2015/19/B/NZ1/00327 (to E.M.D.) from the National Science Centre, Poland, the foundations Carl Tryggers Stiftelse, Magnus Bergvalls Stiftelse and Wilhelm and Martina Lundgrens Vetenskapsfond to MJT, and Stiftelsen Olle Engkvist Byggmästare to NVK, Swedish Research Council VR (grant 637–2014-437) and a Hasselblad Foundation Prize to A. R.

## Conflict of interest statement

None declared.

