## Supplemental information for "The ancillary N-terminal region of the yeast AP-1 transcription factor Yap8 contributes to its DNA binding specificity"

**Content:**

**Supplementary Methods, Tables S1-S4, Figures S1-S5**

**SUPPLEMENTARY METHODS**

**Details of molecular dynamics simulation protocols and analysis procedures**

Molecular dynamics simulations (MD) were carried out with GROMACS MD software package, version 5.1 (1). Simulations were carried out using the combination of the latest AMBER all-atom nucleic acid Parmbsc1 (2) and ff14SB (3) force fields. Each protein-DNA complex was first neutralized with 34 K+ counterions and solvated in a 15Å solvent layer of SCP/E water molecules (4). Additional K+ and Cl- ions were added to reach overall KCl concentration of 150 mM. Each protein-DNA complex was initially energy minimized with 5000 steps of steepest descent, followed by a 500 ps simulation at constant volume, while raising the temperature to 300 K. Afterwards MD simulations were performed at constant pressure and temperature (1 atm., 300 K) using a weak-coupling thermostat (5) with a 0.2 ps coupling constant and an isotropic Parrinello-Rahman barostat (6) with a 2 ps coupling constant, and a 2 fs time step. Bonds involving hydrogen atoms were constrained with the LINCS algorithm (7) and the non-bonded pair list was updated every 20 fs with the group scheme (8). Electrostatic forces were evaluated using particle mesh Ewald algorithm (9) with a real-space cut-off of 10Å. The van der Waals forces were truncated at 10Å also and long-range corrections were added. Center of mass movement was removed every 0.2 ps to eliminate translational kinetic energy (10). Following the initial 100 ns of NPT MD, considered as equilibration, productive runs were recorded for another 0.5 μs for each setup.

**Details of MD analysis procedure**

MD trajectories were analyzed using CPPTRAJ program (11), with a particular focus on the protein-DNA interactions: including hydrogen bonds, salt bridges, and hydrophobic (apolar) interactions. The limit of a direct interaction was set up to be ≤ 3.5Å for a hydrogen bond between relevant heavy atoms, and the angle limit was set up to ≥ 135**°** at the intervening hydrogen atom. While for a salt bridge interaction the limit was set up to 3.5-6.0Å, bearing in mind possible interaction through a bridging water or a bridging counterion. A hydrophobic interaction was defined as a contact ≤ 6Å between the centers of mass of hydrophobic residues (Ala, Ile, Leu, Met, Phe, Trp) and DNA bases. The hydrogen bonds and salt bridges interactions were characterized by a fraction of the trajectory snapshots during which they were maintained, and by the average lifetimes (the lifetime calculations has been smoothed out by ignoring interruption of interactions shorter than 1 ps). Dynamic contacts maps were created by summing up the hydrogen bonds and the salt bridge interactions for each pair of Yap8-DNA interacting resides, which resulted in a contact strength value.

**Supplementary Table S1**

**Table S1.** Plasmids used in this study.

| **Plasmid name** | **Description** | **Reference** |
| --- | --- | --- |
| pYX122 | CEN vector*, HIS3*, *TPI1* promoter |  |
| pYX122-YAP8 | *YAP8-HA* fusion cloned under *TPI1* promoter in pYX122 | (1) |
| pYX122-YAP8-Δ5-13 | *yap8-*Δ*5-13* mutation generated in pYX122-YAP8 | This study |
| pYX122-YAP8-P4A | P4A mutation generated in pYX122-YAP8 | This study |
| pYX122-YAP8-R5A | R5A mutation generated in pYX122-YAP8 | This study |
| pYX122-YAP8-G6A | G6A mutation generated in pYX122-YAP8 | This study |
| pYX122-YAP8-R7A | R7A mutation generated in pYX122-YAP8 | This study |
| pYX122-YAP8-K8A | K8A mutation generated in pYX122-YAP8 | This study |
| pYX122-YAP8-G9A | G9A mutation generated in pYX122-YAP8 | This study |
| pYX122-YAP8-G10A | G10A mutation generated in pYX122-YAP8 | This study |
| pYX122-YAP8-R11A | R11A mutation generated in pYX122-YAP8 | This study |
| pYX122-YAP8-K12A | K12A mutation generated in pYX122-YAP8 | This study |
| pYX122-YAP8-P13A | P13A mutation generated in pYX122-YAP8 | This study |
| pYX122-YAP8-S14A | S14A mutation generated in pYX122-YAP8 | This study |
| pYX122-YAP8-L15A | L15A mutation generated in pYX122-YAP8 | This study |
| pYX122-YAP8-T16A | T16A mutation generated in pYX122-YAP8 | This study |
| pYX122-YAP8-P17A | P17A mutation generated in pYX122-YAP8 | This study |
| pYX122-YAP8-P18A | P18A mutation generated in pYX122-YAP8 | This study |
| pYX122-YAP8-N20A | N20A mutation generated in pYX122-YAP8 | This study |
| pYX122-YAP8-N20Q | N20Q mutation generated in pYX122-YAP8 | This study |
| pYX122-YAP8-N20D | N20D mutation generated in pYX122-YAP8 | This study |
| pYX122-YAP8-K21A | K21A mutation generated in pYX122-YAP8 | This study |
| pYX122-YAP8-Q25A | Q25A mutation generated in pYX122-YAP8 | This study |
| pYX122-YAP8-F33A | F33A mutation generated in pYX122-YAP8 | This study |
| pYX122-YAP8-K35A | K35A mutation generated in pYX122-YAP8 | This study |
| pYX122-YAP8-R36A | R36A mutation generated in pYX122-YAP8 | This study |
| pYX122-YAP8-K37A | K37A mutation generated in pYX122-YAP8 | This study |
| pYX122-YAP8-L38A | L38A mutation generated in pYX122-YAP8 | This study |
| pYX122-YAP8-E39A | E39A mutation generated in pYX122-YAP8 | This study |
| pYX122-YAP8-R40A | R40A mutation generated in pYX122-YAP8 | This study |
| pYX122-YAP8-4aa | Quadruple (A23T L26N S29A N31R) mutation generated in pYX122-YAP8 | This study |
| pYX122-YAP8-7aa | Septuple (A23T L26N S29A N31R K35E L37E E39R) mutation generated in pYX122-YAP8 | This study |
| pYX122-YAP8-8aa | Octuple (N20Q A23T L26N S29A N31R K35E L37E E39R) mutation generated in pYX122-YAP8 | This study |
| pYX122-YAP1 | *YAP1-HA* fusion cloned under *TPI1* promoter in pYX122 | This study |
| pGEX4T-1-GST-YAP8 | *GST-YAP8* fusion for expression in *E. coli* | (2) |
| pGST-YAP8-N20A | N20A mutation generated in pGEX4T-1-GST-YAP8 | This study |
| pGST-YAP8-N20Q | N20Q mutation generated in pGEX4T-1-GST-YAP8 | This study |
| pGST-YAP8-Q25A | Q25A mutation generated in pGEX4T-1-GST-YAP8 | This study |
| pGST-YAP8-4aa | Quadruple (A23T L26N S29A N31R) mutation generated in pGEX4T-1-GST-YAP8 | This study |
| pGST-YAP8-7aa | Septuple (A23T L26N S29A N31R K35E L37E E39R) mutation generated in pGEX4T-1-GST-YAP8 | This study |
| pGST-YAP8-8aa | Octuple (N20Q A23T L26N S29A N31R K35E L37E E39R) mutation generated in pGEX4T-1-GST-YAP8 | This study |
| pGST-YAP8-R7A | R7A mutation generated in pGEX4T-1-GST-YAP8 | This study |
| pGST-YAP8-G10A | G10A mutation generated in pGEX4T-1-GST-YAP8 | This study |
| pGST-YAP8-R11A | R11A mutation generated in pGEX4T-1-GST-YAP8 | This study |
| pSEYC102 | CEN vector, *URA3, lacZ* reporter gene | (3) |
| pEM19 | *ACR3-lacZ* fusion in pSEYC102 | (4) |
| pDP1 | *MUT3-ACR3-lacZ* fusion in pSEYC102 | (2) |

**Supplementary Table S2**

**Table S2.** Oligonucleotides used in this work.

| **Name** | **Sequence 5’ to 3’** |
| --- | --- |
| **Mutagenesis:** | |
| P4A-fw | CATGGGTATGGCAAAAGCGCGTGGAAGAAAAGG |
| P4A-rv | CCTTTTCTTCCACGCGCTTTTGCCATACCCATG |
| R5A-fw | GGTATGGCAAAACCGGCTGGAAGAAAAGGCGGC |
| R5A-rv | GCCGCCTTTTCTTCCAGCCGGTTTTGCCATACC |
| G6A-fw | GTATGGCAAAACCGCGTGCAAGAAAAGGCGGCAGG |
| G6A-rv | CCTGCCGCCTTTTCTTGCACGCGGTTTTGCCATAC |
| R7A-fw | GGCAAAACCGCGTGGAGCAAAAGGCGGCAGGAAG |
| R7A-rv | CTTCCTGCCGCCTTTTGCTCCACGCGGTTTTGCC |
| K8A-fw | CAAAACCGCGTGGAAGAGCAGGCGGCAGGAAGCCTC |
| K8A-rv | GAGGCTTCCTGCCGCCTGCTCTTCCACGCGGTTTTG |
| G9A-fw | CGCGTGGAAGAAAAGCCGGCAGGAAGCCTTCAC |
| G9A-rv | GTGAAGGCTTCCTGCCGGCTTTTCTTCCACGCG |
| G10A-fw | CGTGGAAGAAAAGGCGCCAGGAAGCCTTCACTTAC |
| G10A-rv | GTAAGTGAAGGCTTCCTGGCGCCTTTTCTTCCACG |
| R11A-fw | GGAAGAAAAGGCGGCGCGAAGCCTTCACTTACTC |
| R11A-rv | GAGTAAGTGAAGGCTTCGCGCCGCCTTTTCTTCC |
| K12A-fw | GAAAAGGCGGCAGGGCGCCTTCACTTACTCCAC |
| K12A-rv | GTGGAGTAAGTGAAGGCGCCCTGCCGCCTTTTC |
| P13A-fw | GAAAAGGCGGCAGGAAGGCTTCACTTACTCCACC |
| P13A-rv | GGTGGAGTAAGTGAAGCCTTCCTGCCGCCTTTTC |
| S14A-fw | GGCGGCAGGAAGCCTGCACTTACTCCACCTAAA |
| S14A-rv | TTTAGGTGGAGTAAGTGCAGGCTTCCTGCCGCC |
| L15A-fw | GGCGGCAGGAAGCCTTCAGCTACTCCACCTAAAAA |
| L15A-rv | TTTTTAGGTGGAGTAGCTGAAGGCTTCCTGCCGCC |
| T16A-fw | CAGGAAGCCTTCACTTGCTCCACCTAAAAATAAGAG |
| T16A-rv | CTCTTATTTTTAGGTGGAGCAAGTGAAGGCTTCCTG |
| P17A-fw | GGAAGCCTTCACTTACTGCACCTAAAAATAAGAGAGC |
| P17A-rv | GCTCTCTTATTTTTAGGTGCAGTAAGTGAAGGCTTCC |
| P18A-fw | GCCTTCACTTACTCCAGCTAAAAATAAGAGAGCTGCG |
| P18A-rv | CGCAGCTCTCTTATTTTTAGCTGGAGTAAGTGAAGGC |
| K19A-fw | CCTTCACTTACTCCACCTGCAAATAAGAGACTGCGCAAC |
| K19A-rv | GTTGCGCAGTCTCTTATTTGCAGGTGGAGTAAGTGAAGG |
| N20A-fw | CACTTACTCCACCTAAACAAAAGAGAGCTGCGCAACTTAGAG |
| N20A-rv | CTCTAAGTTGCGCAGCTCTCTTTTGTTTAGGTGGAGTAAGTG |
| N20D-fw | CACTTACTCCACCTAAAGACAAGAGAGCTGCGCAACTTAGAG |
| N20D-rv | CTCTAAGTTGCGCAGCTCTCTTGTCTTTAGGTGGAGTAAGTG |
| N20Q-fw | CACTTACTCCACCTAAACAAAAGAGAACTGCGCAAAATAGAGC |
| N20Q-rv | GCTCTATTTTGCGCAGTTCTCTTTTGTTTAGGTGGAGTAA GTG |
| K21A-fw | CTTACTCCACCTAAAAATGCGAGAGCTGCGCAACTTAGAG |
| K21A-rv | CTCTAAGTTGCGCAGCTCTCGCATTTTTAGGTGGAGTAAG |
| A23T-fw | CCACCTAAAAATAAGAGAACTGCGCAACTTAGAGCATC |
| A23T-rv | GATGCTCTAAGTTGCGCAGTTCTCTTATTTTTAGGTGG |
| Q25A-fw | CTAAAAATAAGAGAGCTGCGGCACTTAGAGCATCCC |
| Q25A-rv | GGGATGCTCTAAGTGCCGCAGCTCTCTTATTTTTAG |
| N31R-fw | CTGCGCAACTTAGAGCATCCCAAAGAGCATTTAGAAAACG |
| N31R-rv | CGTTTTCTAAATGCTCTTTGGGATGCTCTAAGTTGCGCAG |
| F33A-fw | CTTAGAGCATCCCAAAACGCAGCTAGAAAACGAAAGTTGG |
| F33-rv | CCAACTTTCGTTTTCTAGCTGCGTTTTGGGATGCTCTAAG |
| A23TL26NS29A-fw | CCACCTAAAAATAAGAGAACTGCGCAAAATAGAGCAGC |
| A23TL26NS29A-rv | GCTGCTCTATTTTGCGCAGTTCTCTTATTTTTAGGTGG |
| A23TL26NS29AN31R-fw | GAACTGCGCAAAATAGAGCAGCTCAAAGAGCATTTAGAAAACG |
| A23TL26NS29AN31R-rv | CGTTTTCTAAATGCTCTTTGAGCTGCTCTATTTTGCGCAGTTC |
| K35EL37EE39R-fw | CAAAGAGCATTTAGAGAACGAAAGGAGCGAAGATTAGAAGAACTAG |
| K35EL37EE39R-rv | CTAGTTCTTCTAATCTTCGCTCCTTTCGTTCTCTAAATGCTCTTTG |
| K35A-fw | GCATCCCAAAACGCATTTAGAGCACGAAAGTTGGAAAGATTAG |
| K35A-rv | CTAATCTTTCCAACTTTCGTGCTCTAAATGCGTTTTGGGATGC |
| R36A-fw | CCCAAAACGCATTTAGAAAAGCAAAGTTGGAAAGATTAGAAG |
| R36A-rv | CTTCTAATCTTTCCAACTTTGCTTTTCTAAATGCGTTTTGGG |
| K37A-fw | CCCAAAACGCATTTAGAAAACGAGCGTTGGAAAGATTAGAAG |
| K37A-rv | CTTCTAATCTTTCCAACGCTCGTTTTCTAAATGCGTTTTGGG |
| L38A-fw | CGCATTTAGAAAACGAAAGGCGGAAAGATTAGAAGAACTAGAG |
| L38A-rv | CTCTAGTTCTTCTAATCTTTCCGCCTTTCGTTTTCTAAATGCG |
| E39A-fw | GCATTTAGAAAACGAAAGTTGGCAAGATTAGAAGAACTAGAGAAG |
| E39A-rv | CTTCTCTAGTTCTTCTAATCTTGCCAACTTTCGTTTTCTAAATGC |
| R40A-fw | GAAAACGAAAGTTGGAAGCATTAGAAGAACTAGAGAAGAAG |
| R40A-rv | CTTCTTCTCTAGTTCTTCTAATGCTTCCAACTTTCGTTTTC |
| **qRT-PCR:** | |
| ACR3-fw | CGGCATACCACTGGGAATT |
| ACR3-rv | GCACCAATGGGACAAAGCA |
| PRACR3-fw | TTACGCTTGCTGGATTGTCA |
| PRACR3-rv | CGTTGCCGCTAAAGTTGATT |
| IPP1-fw | CTTTATTGGATGAAGGTGA |
| IPP1-rv | TTAATTGTTTCCAGGAGTC |
| **EMSA:** | |
| ACR3short-fw | (Btn)TCTTAATTATCTTTTTGTTTGATTAATAATCAACTTTAGCGGCAACGCTCC |
| ACR3short-rv | (Btn)GGAGCGTTGCCGCTAAAGTTGATTATTAATCAAACAAAAAGATAATTAAGA |
| MUT3short-fw | TCTTAATTATCTTTTTGTTTGATTACTAATCAACTTTAGCGGCAACGCTCC |
| MUT3short-rv | GGAGCGTTGCCGCTAAAGTTGATTAGTAATCAAACAAAAAGATAATTAAGA |
| TRX2short-fw | ATTGTTTATACTCTTAGTAAAGGATGCTCCCTACAAGGTGGCTCTTTTCTTACTAAGCGCGTTCAGTTTC |
| TRX2short-rv | GAAACTGAACGCGCTTAGTAAGAAAAGAGCCACCTTGTAGGGAGCATCCTTTACTAAGAGTATAAACAAT |
| GSH1short-fw | TTCTGCCCAACGACGGCTGCCATTAGTCAGCATGGCGCGCACGTGACTACA |
| GSH1short-rv | TGTAGTCACGTGCGCGCCATGCTGACTAATGGCAGCCGTCGTTGGGCAGAA |

**Supplementary Table S3**

Table S3. Interactions between Yap8 wild-type protein dimer and DNA characterized by the percentage presence during 0.5 μs MD simulation and the average lifetime (<LT>)(ps). Interactions in orange represent the salt bridges, in black – the hydrogen bonds between the DNA backbone and the protein, and in blue – the hydrogen bonds between the DNA bases and the protein. The table is limited to the interactions that occur at least 5% of the time of the MD simulation.

| **Contact name** | **%** | **<LT> (ps)** | **Contact name** | **%** | **<LT> (ps)** |
| --- | --- | --- | --- | --- | --- |
| **Yap8 wild-type monomer 1** | | | | | |
| T6_w_ (OP2)-Arg11 (N) | 99 | 2157 | T2_w_ (O2)-Arg7 (NH1) | 26 | 15 |
| T10_w_ (OP1)-Arg36 (NH1) | 99 | 868 | A3_c_ (OP2)-Arg11 (NH2) | 25 | 196 |
| A9_w_ (OP1)-Ser29 (OG) | 94 | 564 | T7_w_ (OP2)-Arg11 (NE) | 19 | 52 |
| T6_w_ (OP2)-Lys12 (N) | 93 | 612 | A14_c_ (OP1)-Arg34 (NH2) | 16 | 22 |
| T3_w_ (O2)-Arg7 (NH2) | 91 | 1006 | T12_c_ (O5')-Arg27 (NH1) | 15 | 27 |
| T5_w_ (O3')-Gly10 (N) | 87 | 501 | G8_w_ (O5')-Asn25 (NE2) | 14 | 13 |
| A1_c_ (OP2)-Lys8 (N) | 81 | 82 | G8_w_ (OP2)-Asn25 (NE2) | 14 | 12 |
| T5_w_ (OP2)-Arg7 (N) | 72 | 66 | T7_w_ (OP2)-Arg11 (NH1) | 14 | 26 |
| A11_c_ (N6)-Gln30 (OE1) | 72 | 53 | T12_c_ (OP1)-Arg27 (NE) | 14 | 30 |
| A2_c_ (O3')-Lys8 (N) | 69 | 34 | T7_w_ (OP1)-Arg22 (NH2) | 14 | 22 |
| G8_w_ (O6)-Arg22 (NH1) | 51 | 71 | A1_c_ (OP1)-Lys8 (NZ) | 13 | 13 |
| A3_c_ (OP2)-Arg11 (NH1) | 50 | 38 | T13_c_ (OP1)-Arg34 (NH2) | 11 | 40 |
| G8_w_ (N7)-Arg22 (NH2) | 49 | 94 | T5_w_ (OP2)-Lys12 (NZ) | 10 | 23 |
| T7_w_ (OP2)-Arg11 (NH2) | 44 | 79 | G8_w_ (N7)-Arg22 (NH1) | 9 | 13 |
| G8_w_ (OP1)-Asn25 (NE2) | 43 | 22 | T13_c_ (OP1)-Arg34 (NH1) | 9 | 27 |
| T3_w_ (O2)-Arg7 (NH1) | 40 | 17 | T7_w_ (OP2)-Arg22 (NH1) | 8 | 13 |
| T12_c_ (OP1)-Arg27 (NH1) | 38 | 30 | T3_w_ (O4')-Arg7 (NH1) | 8 | 8 |
| T7_w_ (OP1)-Arg22 (NH1) | 36 | 92 | G8_w_ (O6)-Arg22 (NH2) | 7 | 16 |
| A3_c_ (OP2)-Arg11 (NE) | 33 | 166 | T12_c_ (O5')-Arg27 (NH2) | 7 | 14 |
| T10_w_ (O4)-Gln30 (NE2) | 33 | 96 | T6_w_ (O3')-Arg11 (NH2) | 7 | 20 |
| A14_c_ (OP1)-Arg34 (NH1) | 31 | 48 | T7_w_ (OP1)-Arg11 (NE) | 7 | 20 |
| A11_c_ (N6)-Gln30 (NE2) | 29 | 20 | T7_w_ (OP1)-Arg11 (NH2) | 6 | 48 |
| T6_w_ (OP1)-Lys12 (NZ) | 28 | 51 | A11_c_ (N7)-Gln30 (NE2) | 6 | 27 |
| T12_c_ (OP1)-Arg27 (NH2) | 28 | 81 | A11_c_ (OP1)-Arg27 (NH1) | 6 | 53 |
| C4_c_ (O3')-Arg11 (NH1) | 28 | 24 | A14_c_ (O5')-Arg34 (NH2) | 5 | 11 |
| **Yap8 wild-type monomer 2** | | | | | |
| T15_c_ (O4)-Gln30 (NE2) | 100 | 922 | A13_w_ (N6)-Arg34 (NH2) | 14 | 18 |
| G18_c_ (O6)-Arg22 (NH1) | 91 | 154 | A17_c_ (OP1)-Ser29 (OG) | 14 | 33 |
| T19_c_ (OP1)-Lys21 (NZ) | 82 | 90 | T19_c_ (OP2)-Arg11 (NE) | 12 | 41 |
| T14_w_ (OP1)-Arg27 (NE) | 80 | 694 | A13_w_ (OP2)-Arg27 (NE) | 11 | 30 |
| T14_w_ (OP1)-Arg27 (NH1) | 78 | 172 | T24_w_ (OP1)-Lys8 (NZ) | 11 | 13 |
| T24_w_ (OP2)-Lys8 (N) | 60 | 64 | A12_w_ (O5')-Arg34 (NH2) | 11 | 13 |
| G18_c_ (N7)-Arg22 (NH2) | 56 | 29 | T24_w_ (OP1)-Lys8 (N) | 11 | 38 |
| G18_c_ (O6)-Arg22 (NH2) | 54 | 27 | T19_c_ (OP2)-Arg11 (N) | 10 | 263 |
| A17_c_ (OP1)-Arg36 (NH2) | 54 | 58 | A25_w_ (OP2)-Arg7 (NH1) | 10 | 26 |
| T16_c_ (OP1)-Arg36 (NH1) | 53 | 56 | T23_w_ (OP1)-Lys8 (NZ) | 9 | 20 |
| T22_w_ (O2)-Arg7 (NH1) | 50 | 202 | T23_w_ (OP2)-Arg11 (NE) | 9 | 42 |
| A12_w_ (OP1)-Arg34 (NH1) | 50 | 135 | T23_w_ (OP2)-Arg11 (NH1) | 9 | 22 |
| T19_c_ (OP2)-Ser14 (OG) | 43 | 169 | T23_w_ (OP2)-Lys8 (NZ) | 9 | 16 |
| T23_w_ (O3')-Lys8 (N) | 40 | 28 | A25_w_ (OP1)-Arg7 (NE) | 8 | 26 |
| T14_w_ (O4)-Gln30 (NE2) | 37 | 21 | A13_w_ (O5')-Arg27 (NE) | 8 | 15 |
| T19_c_ (OP1)-Asn25 (NE2) | 35 | 75 | A22_c_ (N3)-Arg7 (NH2) | 7 | 53 |
| A13_w_ (N7)-Arg34 (NH2) | 34 | 69 | A23_c_ (N3)-Arg7 (NH2) | 7 | 601 |
| T14_w_ (OP2)-Arg27 (NH1) | 33 | 12 | A13_w_ (OP1)-Arg27 (NE) | 7 | 15 |
| T22_w_ (O2)-Arg7 (NE) | 32 | 48 | T24_w_ (OP1)-Gly9 (N) | 7 | 50 |
| A13_w_ (OP1)-Asn31 (ND2) | 30 | 67 | A15_w_ (OP1)-Lys19 (NZ) | 6 | 25 |
| A17_c_ (OP1)-Arg36 (NH1) | 25 | 39 | T23_w_ (OP2)-Arg11 (NH2) | 6 | 28 |
| T19_c_ (O5')-Asn25 (NE2) | 25 | 23 | T20_c_ (O3')-Ser14 (OG) | 6 | 12 |
| A12_w_ (OP1)-Arg34 (NH2) | 24 | 36 | T23_w_ (OP1)-Arg11 (NH1) | 6 | 25 |
| G18_c_ (OP1)-Asn25 (NE2) | 23 | 119 | A23_c_ (O4')-Arg7 (NH1) | 6 | 41 |
| A12_w_ (OP1)-Arg34 (NE) | 21 | 30 | G18_c_ (N7)-Arg22 (NH1) | 6 | 25 |
| T19_c_ (OP2)-Arg11 (NH1) | 21 | 52 | T19_c_ (O5')-Arg11 (NH1) | 6 | 20 |
| A13_w_ (OP2)-Arg27 (NH1) | 17 | 200 | A25_w_ (OP1)-Arg7 (NH1) | 5 | 21 |
| A13_w_ (OP1)-Arg34 (NH1) | 15 | 32 | T16_c_ (OP1)-Arg36 (NE) | 5 | 11 |
| T16_c_ (OP1)-Arg36 (NH2) | 15 | 155 |  |  |  |

Supplementary Table S4

Table S4. Interactions between Yap8 Asn20Ala mutant protein dimer and DNA characterized by the percentage presence during 0.5 μs MD simulation and the average lifetime (<LT>)(ps). Interactions in orange represent the salt bridges, in black – the hydrogen bonds between the DNA backbone and the protein, and in blue – the hydrogen bonds between the DNA bases and the protein. The table is limited to the interactions that occur at least 5% of the time of the MD simulation.

| **Contact name** | **%** | **<LT> (ps)** | **Contact name** | **%** | **<LT> (ps)** |
| --- | --- | --- | --- | --- | --- |
| **Yap8 Asn20Ala mutant monomer 1** | | | | | |
| A9_w_ (OP1)-Ser29 (OG) | 100 | 2080 | A11_c_ (N6)-Gln30 (NE2) | 20 | 12 |
| T10_w_ (OP1)-Arg36 (NH1) | 99 | 737 | A11_c_ (N6)-Gln30 (NE2) | 20 | 12 |
| T10_w_ (O4)-Gln30 (NE2) | 98 | 1018 | T12_c_ (O5')-Arg27 (NH2) | 20 | 16 |
| T3_w_ (O2)-Arg7 (NH2) | 89 | 194 | T2_w_ (O2)-Arg7 (NH1) | 16 | 14 |
| T7_w_ (OP1)-Arg22 (NH1) | 83 | 144 | T5_w_ (O3')-Lys12 (NZ) | 14 | 15 |
| T12_c_ (OP1)-Arg27 (NH2) | 75 | 80 | T6_w_ (OP1)-Lys12 (NZ) | 14 | 83 |
| T12_c_ (OP1)-Arg27 (NH1) | 68 | 53 | A2_c_ (O3')-Gly9 (N) | 14 | 15 |
| T7_w_ (OP1)-Arg22 (NE) | 66 | 36 | A2_c_ (O3')-Lys8 (N) | 13 | 13 |
| A1_c_ (OP2)-Lys8 (N) | 65 | 80 | A2_c_ (OP2)-Arg11 (N) | 12 | 115 |
| T3_w_ (O2)-Arg7 (NH1) | 59 | 25 | T5_w_ (OP2)-Lys12 (NZ) | 11 | 41 |
| T5_w_ (OP2)-Arg7 (N) | 54 | 68 | A2_c_ (OP2)-Lys12 (NZ) | 10 | 31 |
| A11_c_ (N6)-Gln30 (OE1) | 48 | 22 | A1_c_ (OP1)-Lys8 (NZ) | 9 | 14 |
| A14_c_ (OP1)-Arg34 (NH1) | 45 | 35 | A3_c_ (OP2)-Lys12 (NZ) | 8 | 87 |
| G8_w_ (O5')-Gln25 (NE2) | 45 | 33 | T13_c_ (OP1)-Arg34 (NH2) | 7 | 49 |
| T7_w_ (O5')-Arg22 (NE) | 38 | 16 | C4_c_ (O2)-Lys12 (NZ) | 7 | 71 |
| A14_c_ (OP1)-Arg34 (NH2) | 31 | 45 | T6_w_ (OP1)-Arg22 (NH1) | 7 | 21 |
| A1_c_ (OP2)-Gly9 (N) | 28 | 45 | T3_w_ (O4')-Arg7 (NH1) | 6 | 8 |
| T6_w_ (OP2)-Lys12 (NZ) | 28 | 34 | A11_c_ (OP1)-Arg27 (NH1) | 6 | 55 |
| G8_w_ (OP1)-Gln25 (NE2) | 27 | 25 | T6_w_ (OP2)-Arg22 (NH1) | 5 | 22 |
| T12_c_ (O5')-Arg27 (NH1) | 24 | 23 | A1_c_ (O3')-Arg7 (NE) | 5 | 25 |
| **Yap8 Asn20Ala mutant monomer 2** | | | | | |
| T15_c_ (O4)-Gln30 (NE2) | 100 | 1187 | A16_w_ (N7)-Arg22 (NH1) | 13 | 22 |
| A12_w_ (OP1)-Arg34 (NH2) | 91 | 317 | T22_w_ (O2)-Arg7 (NH1) | 13 | 64 |
| A17_c_ (OP1)-Ser29 (OG) | 89 | 740 | T20_c_ (OP2)-Arg7 (NH1) | 12 | 40 |
| T16_c_ (OP1)-Arg36 (NH1) | 87 | 1810 | T19_c_ (OP2)-Lys21 (NZ) | 12 | 11 |
| T19_c_ (OP1)-Gln25 (NE2) | 85 | 46 | T20_c_ (OP2)-Lys12 (NZ) | 12 | 27 |
| T19_c_ (O5')-Gln25 (NE2) | 80 | 37 | A13_w_ (O5')-Arg27 (NE) | 12 | 14 |
| A12_w_ (OP1)-Arg34 (NH1) | 80 | 83 | T20_c_ (OP2)-Arg11 (NH2) | 10 | 58 |
| T14_w_ (OP1)-Arg27 (NE) | 66 | 550 | A17_c_ (N7)-Ser29 (OG) | 10 | 104 |
| G18_c_ (O6)-Arg22 (NH1) | 63 | 52 | T17_w_ (O4)-Arg22 (NH1) | 8 | 12 |
| T14_w_ (OP1)-Arg27 (NH1) | 63 | 123 | T24_w_ (OP1)-Lys8 (NZ) | 8 | 23 |
| T24_w_ (OP2)-Lys8 (N) | 60 | 172 | T23_w_ (OP1)-Lys8 (NZ) | 8 | 14 |
| T19_c_ (OP1)-Lys21 (NZ) | 57 | 32 | T17_w_ (O4)-Arg22 (NH2) | 7 | 137 |
| T14_w_ (O4)-Gln30 (NE2) | 56 | 22 | T20_c_ (OP2)-Arg7 (NH2) | 7 | 46 |
| G18_c_ (O6)-Arg22 (NE) | 55 | 85 | T20_c_ (OP2)-Arg11 (NH1) | 7 | 42 |
| A13_w_ (OP1)-Asn31 (ND2) | 52 | 75 | T16_c_ (OP1)-Arg36 (NH2) | 7 | 89 |
| T24_w_ (OP2)-Arg7 (NH2) | 29 | 347 | T23_w_ (O3')-Lys8 (N) | 7 | 27 |
| A13_w_ (OP2)-Arg27 (NH1) | 29 | 159 | A17_c_ (O5')-Ser29 (OG) | 7 | 7 |
| A12_w_ (O5')-Arg34 (NH1) | 25 | 11 | A25_w_ (OP2)-Arg7 (NE) | 6 | 47 |
| T23_w_ (OP2)-Lys8 (NZ) | 24 | 16 | T24_w_ (OP2)-Arg7 (NE) | 6 | 57 |
| A25_w_ (N3)-Arg7 (NH2) | 23 | 82 | G18_c_ (O3')-Ser29 (OG) | 6 | 6 |
| T14_w_ (OP2)-Arg27 (NH1) | 23 | 11 | A13_w_ (N7)-Arg34 (NH1) | 6 | 33 |
| G18_c_ (O6)-Arg22 (NH2) | 23 | 104 | A25_w_ (N3)-Arg7 (NE) | 5 | 50 |
| A13_w_ (OP1)-Arg27 (NE) | 19 | 23 | T24_w_ (OP1)-Lys8 (N) | 5 | 22 |
| A13_w_ (OP2)-Arg27 (NE) | 16 | 26 |  |  |  |

**Supplementary Figure S1**

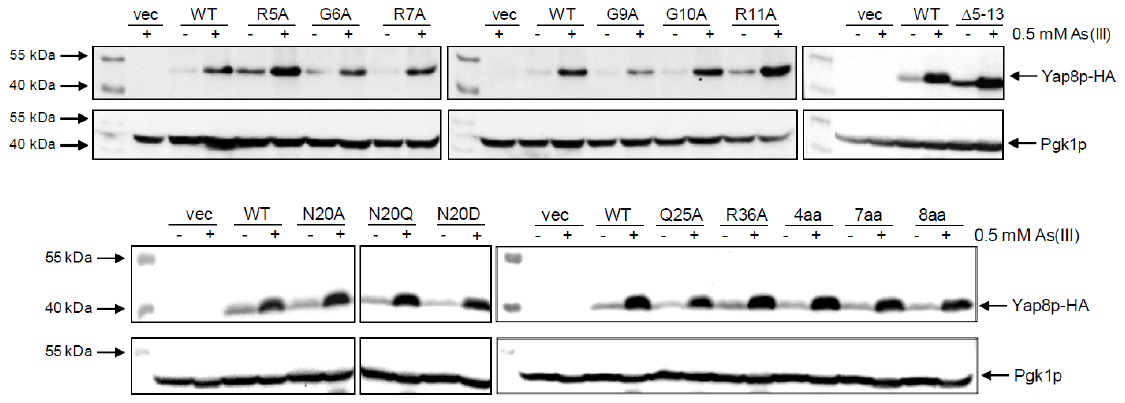

**Figure S1.** Protein level of Yap8 variants. Western blot analysis of total protein extracts prepared from the *yap8*Δ mutant transformed with the plasmids expressing indicated variants of Yap8-HA fusion protein under the control of constitutive *TPI1* promoter. Proteins were isolated from cells cultivated in standard conditions (control) or exposed to 0.5 mM As(III) for 30 min. The anti-HA antibodies were used to detect Yap8. Levels of 3-phosphoglycerate kinase Pgk1 detected with the anti-PGK1 antibodies served as a loading control.

**Supplementary Figure S2**

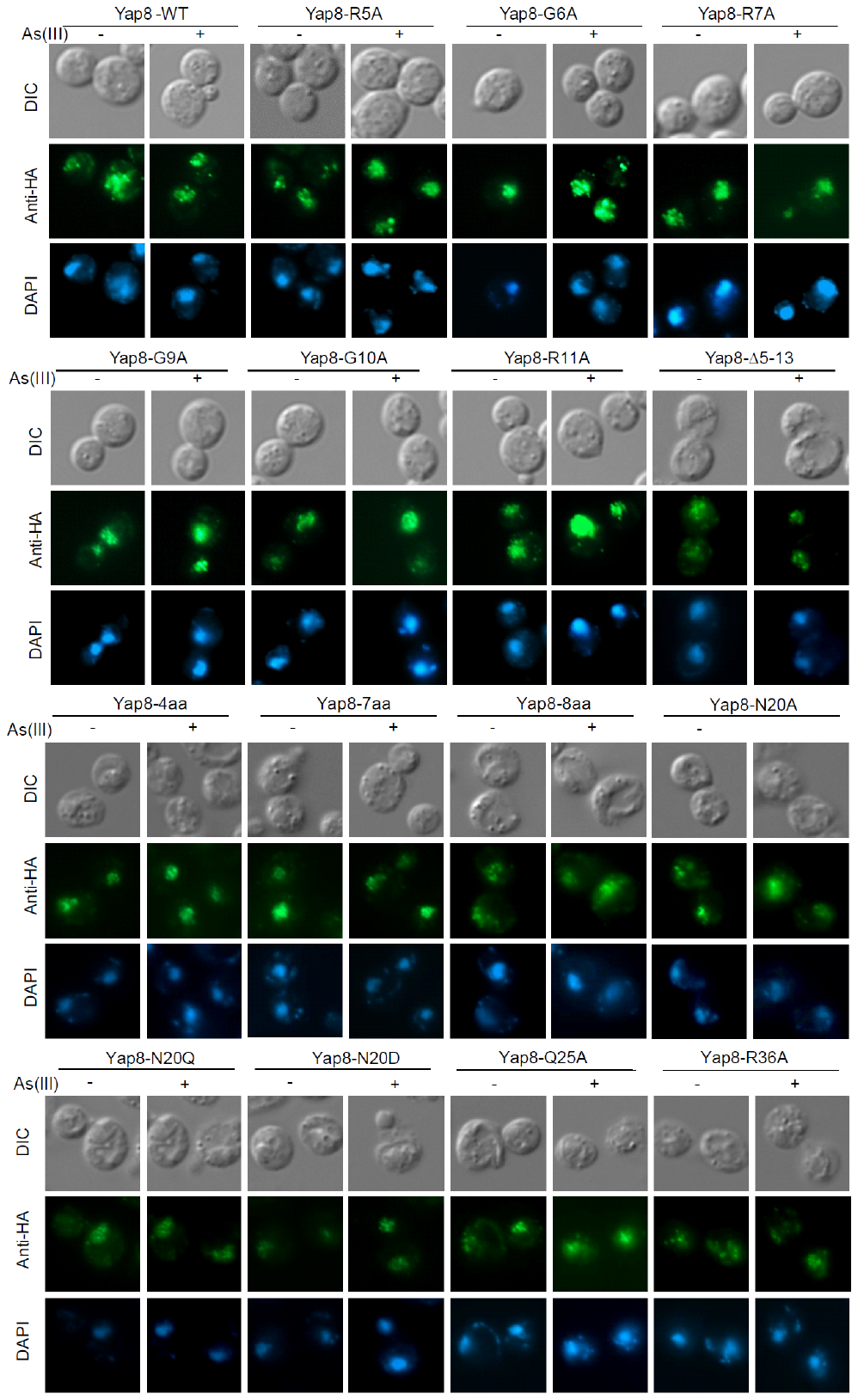

**Figure S2.** Subcellular localization of Yap8 variants. The *yap8*Δ mutant was transformed with pYX122-based plasmids expressing indicated mutant variants of Yap8-HA fusion protein under the control of constitutive *TPI1* promoter. Cells were cultivated in standard conditions (control) or exposed to 0.5 mM As(III) for 30 min and then subjected to immunofluorescence microscopy using primary anti-HA antibody and secondary Alexa Fluor® 488-labeled antibody to detect subcellular localization of mutant variants of Yap8-HA. Cells were also stained with DAPI to visualize nuclei and analyzed by differential interference contrast (DIC)*.*

**Supplementary Figure S3**

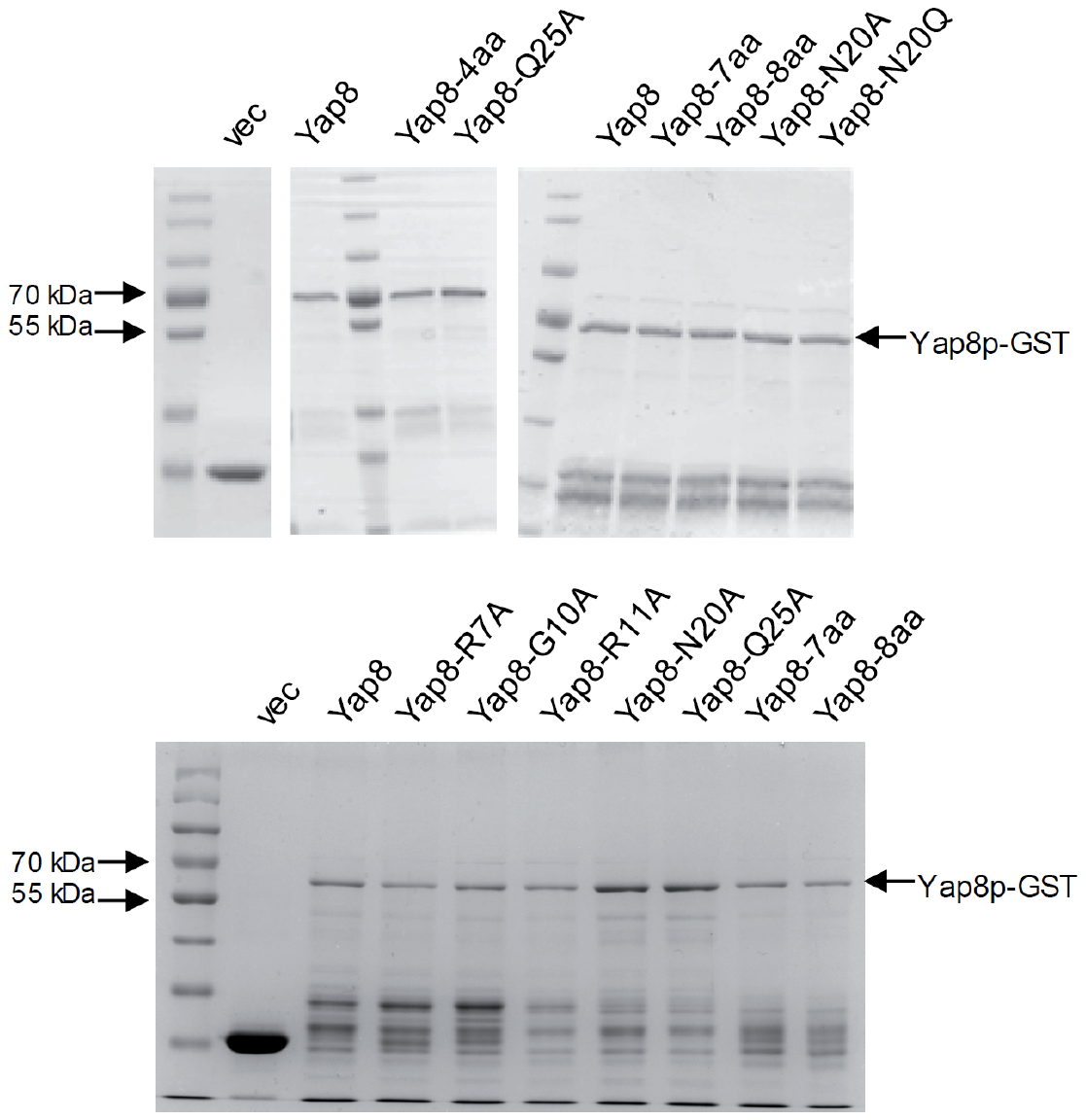

**Figure S3.** SDS-PAGE analysis of purified variants of GST-YAP8 proteins*.* Protein extracts of wild type and indicated mutant variants of GST-YAP8 fusion were isolated from *E. coli* and purified using glutathione sepharose affinity chromatography resin (GE Healthcare Life Sciences). Samples (∼10 ng per lane) were loaded on a 10% polyacrylamide gel, subjected to SDS-PAGE and stained with Coomassie blue. From left: lane 1) Protein Marker, (PageRuler™ Prestained Protein Ladder, Thermo Scientific); lane 2) empty vector with GST; lanes 3-10) variants of GST-YAP8. The upper panel refers to Figure 4A, the lower panel refers to Figure 3A and C, Figure 6A and C.

**Supplementary Figure S4**

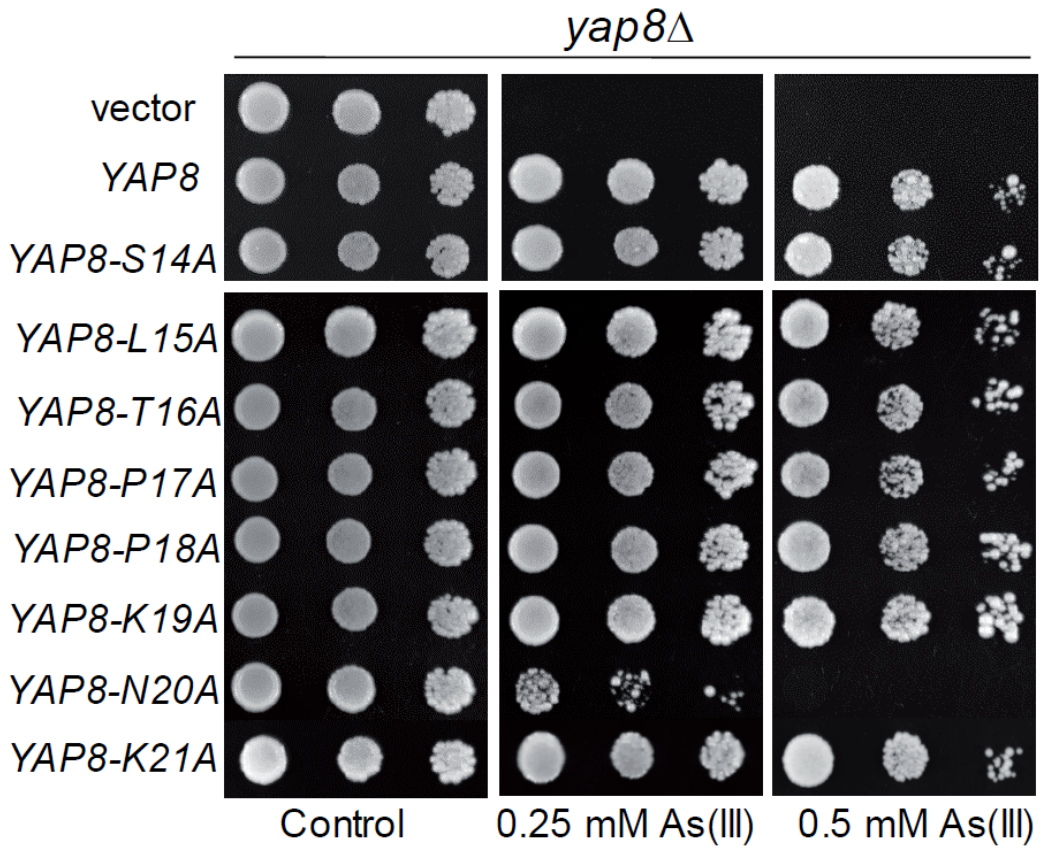

**Figure S4.** Mutational analysis of the N-terminal region adjacent to the basic region of Yap8. The *yap8*Δ mutant was transformed with empty vector (pYX122) or plasmids expressing indicated Yap8 variants. The resulting transformants were spotted on minimal selective plates containing various concentrations of As(III) and incubated 3 days at 28°C.

**Supplementary Figure S5**

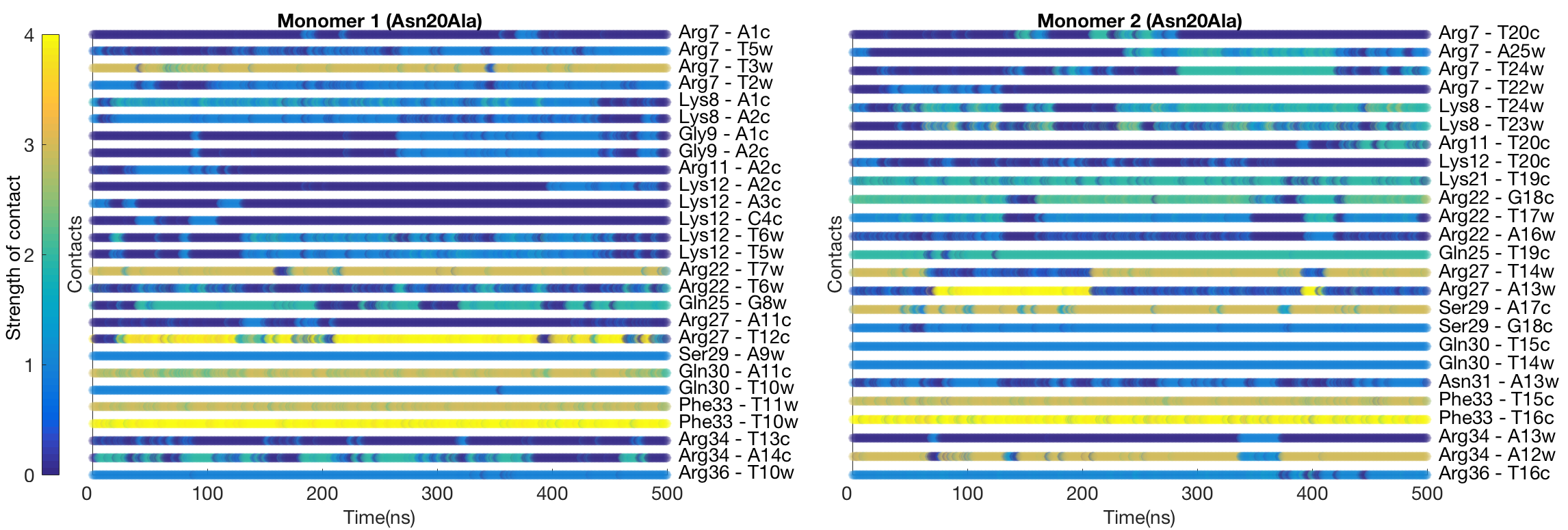

Figure S5. Dynamic interactions maps illustrating the intermolecular DNA-Yap8 (N20A) mutant interface. The interactions between pairs of the protein-DNA residues are characterized by a contact strength and its occurrence during the 0.5 μs MD simulation.
